# Topologically-based parameter inference for agent-based model selection from spatiotemporal cellular data

**DOI:** 10.1101/2025.06.13.659586

**Authors:** Alyssa R. Wenzel, Patrick M. Haughey, Kyle C. Nguyen, John T. Nardini, Jason M. Haugh, Kevin B. Flores

## Abstract

Advances in spatiotemporal single-cell imaging have enabled detailed observations of cell population dynamics and intercellular interactions. However, translating these rich data sets into mechanistic insight remains a significant challenge. Agent-based models (ABMs) are a bottom-up computational framework for investigating the emergent behavior of cell populations that can arise from rules defining the interactions between individual neighboring cells, while topological data analysis (TDA) provides robust descriptors of spatial organization. We present TOPAZ (TOpologically-based Parameter inference for Agent-based model optimiZation), a computational pipeline that integrates TDA with approximate Bayesian computation (ABC), approximate approximate Bayesian computation (AABC), and Bayesian model selection to identify biologically plausible ABMs from spatiotemporal cellular data. TOPAZ uses persistent homology to quantify spatial features of cell trajectories and combines this topological information with parameter inference via ABC and AABC and model comparison using the Bayesian information criterion. We validate TOPAZ using simulations of collective fibroblast movement, demonstrating its ability to accurately recover model parameters and distinguish between a baseline ABM and an extended model that incorporates an alignment interaction. Our results and open-source code demonstrate the utility of TOPAZ as an extensible framework for mechanistic inference and model discrimination in spatial single-cell analysis.

**Author summary:** Understanding how individual cells coordinate to produce complex collective behaviors is a major challenge in computational biology, especially with the increasing availability of high-resolution, spatiotemporal single-cell data. While agent-based models (ABMs) offer a flexible framework for simulating cell behaviors and interactions, they are often difficult to calibrate and compare. Topological data analysis (TDA), on the other hand, captures spatial organization in a robust and scale-invariant way but lacks mechanistic interpretability. In this work, we present TOPAZ (TOpologically-based Parameter inference for Agent-based model optimiZation), a novel computational pipeline that integrates TDA with approximate Bayesian computation, approximate approximate Bayesian computation, and Bayesian model selection to infer biologically meaningful parameters and identify the most plausible ABM from spatiotemporal cellular data. We benchmark TOPAZ using synthetic data from ABMs of collective cell movement in dense fibroblast populations. Our results show that TOPAZ can distinguish between competing mechanistic hypotheses, namely the presence or absence of alignment interactions among neighboring cells. This approach provides a powerful and extensible framework for model inference and selection with the potential to enable deeper insights into the mechanisms driving complex emergent behaviors in cell populations.

## Introduction

Advances in microscopy imaging and live-cell tracking have enabled the high-throughput collection of single-cell spatiotemporal dynamics across a wide range of biological systems [1, 2]. A central challenge in computational biology is to develop integrative methods that extract mechanistic insight from multi-scale datasets, particularly linking subcellular signaling dynamics to population-level behavior, such as collective cell migration at the tissue level. Connecting these scales with computational inference and mechanistic modeling has the potential to provide data-driven insight into processes that are fundamental to morphogenesis, wound healing, and cancer progression [3]. Agent-based models (ABMs) are computational experiments that model individual agents and how they interact with other agents and the environment [4, 5]. They have recently been used to computationally investigate the mechanisms through which cell-to-cell heterogeneity, derived from single-cell transcriptomics and proteomics data, gives rise to population-level behaviors, e.g., by leveraging single-cell data for ABM initialization and calibration [6]. Topological data analysis (TDA) is another computational framework that has been utilized in bioinformatics studies for analyzing single-cell omics data. It uses concepts from topology to describe the shape of data and can be used to detect changes between distinct simulations that vary over time and space [7]. For example, TDA has been used to infer the geometric structure of cellular niches and spatial expression gradients by using persistent homology to identify spatially coherent gene expression domains and to characterize tissue architecture in high-dimensional spatial omics datasets [8, 9]. Yet, while TDA excels at quantifying structural organization features in spatial single-cell data, such as gradients, boundaries, and connectivity, it is not inherently mechanistic and does not capture causal or dynamical processes. Conversely, ABMs provide a bottom-up framework for simulating individual cell behaviors and interactions, but are computationally intensive, sensitive to parameter choices, and often difficult to fit or validate directly against data. This motivates the development of hybrid approaches that integrate TDA and ABMs to bridge intracellular, intercellular, and population-level scales to enable mechanistic inference grounded in topological summaries of spatial omics data.

Previous work exemplified that TDA can be used for extracting multi-scale structural features from high-dimensional data, which can be used to inform dynamical modeling frameworks such as ABMs [10, 11]. While these studies demonstrate the utility of TDA in extracting structural features, they primarily offer descriptive insights without establishing direct links between topological summaries and specific mechanistic parameters within dynamical models. This gap was addressed in part by Nguyen et al., who developed a framework combining TDA with Approximate Bayesian Computation (ABC) for parameter inference in an ABM of collective fibroblast motion [12]. Nguyen et al. demonstrated how persistent homology can be applied to single-cell trajectory data to extract summary statistic for performing likelihood-free inference with ABC.

However, the model considered in their study considered a single ABM that only included repulsion and attraction forces and did not consider additional biologically motivated mechanisms such as directional alignment or anisotropy [13, 14]. Moreover, while Nguyen et al. exemplified that parameter inference with TDA was possible, the biologically-motivated mechanisms lacked a model selection component that could formally distinguish between competing hypotheses of cellular interaction. These shortcomings highlight the need for a unified data-driven pipeline that couples spatiotemporal single-cell data, TDA, ABMs and statistical model comparison for developing mechanistically interpretable and data-constrained simulations of cellular population dynamics. Specifically, model comparison is a step often overlooked in ABM pipelines which is essential to ensure that model complexity is justified by explanatory power [15, 16]. We demonstrate how integrating model selection into the TDA-ABC framework yields a novel and extensible approach for analyzing complex collective cell behaviors in biomedical contexts and how adding a directional alignment mechanism may lead to more biologically-accurate results.

### Agent-based models

In this work, we focus on ABMs of single-cell dynamics informed by time-lapse imaging data. Specifically, we investigate how simple interaction rules can give rise to emergent collective behaviors, such as the parallel (and anti-parallel) streaming—or *fluidization*—observed in dense fibroblast monolayers. To capture these dynamics, we adopt the D’Orsogna ABM [12, 17]. We use the D’Orsogna model, referred to here as Model_DO_, as an example of an ABM for which minimal interaction rules can give rise to many distinct population-level movement patterns, some of which recapitulate collective cell migration. Model_DO_ describes the dynamics of self-propelled agents with uniform mass, subject to linear drag and pairwise interactions governed by attractive and repulsive forces, with magnitudes *C*_*a*_ and *C*_*r*_, and length scales *ℓ*_*a*_ and *ℓ*_*r*_. These parameters capture biologically motivated behaviors such as contact inhibition of locomotion (repulsion) and matrix-mediated long-range attraction, as described in [18–20]. A full description of the Model_DO_ ABM can be found in Supplementary Material Appendix S1.

To assess whether distinct mechanisms of interaction can be reliably distinguished using summary statistics derived from TDA, we compare two ABMs: the baseline D’Orsogna model (Model_DO_) and an extended model with alignment interactions (Model_AL_). We created Model_AL_ to determine if the addition of alignment into Model_DO_ leads to the emergence of fluidization, as seen in experiments of fibroblast migration in [12]. We simulate synthetic trajectory data from each model across a range of parameter settings and then fit the alternative model to that data using ABC and AABC. By applying model selection via the Bayesian information criterion (BIC), we assess whether the inference pipeline can correctly recover the true generative model. This simulation-based framework serves as a proof-of-concept for using TDA-informed ABC and AABC to distinguish between competing mechanistic hypotheses—a critical step toward future applications involving real spatial single-cell datasets that incorporate omics-level molecular information. For example, Johnson et al. incorporate spatial transcriptomic sequencing of resected pancreatic lesions to initialize agent-based models, noting that these high-resolution omics datasets may also enable systematic model selection among competing mechanistic hypotheses [21].

We nondimensionalize the D’Orsogna model and express the dynamics in terms of two key parameters: C = *C*_*r*_*/C*_*a*_ and L = *ℓ*_*r*_*/ℓ*_*a*_, representing relative attraction strength and interaction range, respectively. These parameters are sufficient to reproduce a variety of collective behaviors in self-propelled cell populations [12].

To capture alignment effects observed in fibroblasts, we extend the model to include a third parameter W, which governs the strength of orientation coupling (see Supplementary Material Appendix S1 for a derivation from Model_DO_). This extension is biologically motivated by experimental observations of contact-mediated alignment [22] and theoretical models of directional organization in cell populations [23]. In our extended model (Model_AL_), agents adjust their velocity angle based on local alignment while retaining the original D’Orsogna speed dynamics. Thus, the baseline model estimates two parameters (C, L), while the alignment model estimates three (C, L, W). Although our motivating application is fibroblast migration, the models are general and relevant to a range of active biological systems governed by short- and long-range interactions.

### TOpologically-based Parameter inference for Agent-based model optimiZation (TOPAZ)

We developed a computational pipeline named “TOPAZ” that combines TDA computed from ABM simulations, ABC, approximate approximate Bayesian computation (AABC), and BIC for ABM model selection derived from spatiotemporal cellular data. Fig 1 displays the computational steps in the pipeline. **Step 1:** TDA attributes in the form of “Contour Realization Of Computed K-dimensional hole Evolution in the Rips complex” (Crocker) [24] plots are computed using the Vietoris-Rips filtration from spatiotemporal point clouds, in the form (*x, y, θ*), derived from experimental data or ABM simulations. **Step 1a:** Visualization with dimensionality reduction techniques, such as t-distributed stochastic neighbor embedding (t-SNE) or principal component analysis, are recommended. **Step 2:** TDA diagrams are used with ABC as summary statistics for parameter inference and posterior density generation as in [12]. **Step 2a:** Use of AABC to generate more samples for parameter inference if needed. **Step 2b:** Assessment of whether posterior predictive summary statistic distributions are statistically distinguishable across candidate models using Permutational Multivariate Analysis of Variance (PERMANOVA), Energy Distance, and/or Maximum Mean Discrepancy (MMD) prior to BIC-based model selection. For more details about statistical tests before BIC, see S3 Appendix. **Step 3:** The posterior distribution from the top 1% of ABC samples, the number of parameters, and the number of data points are used to calculate the BIC score for the ABM. **Step 4:** The ABM with the lowest BIC score is selected as the optimal model. We use BIC instead of other information criterion, such as Akaike information criterion (AIC), due to the log-likelihood of the sum of squared errors being so large and our desire to emphasize the penalty for the number of parameters.

**Fig 1.**
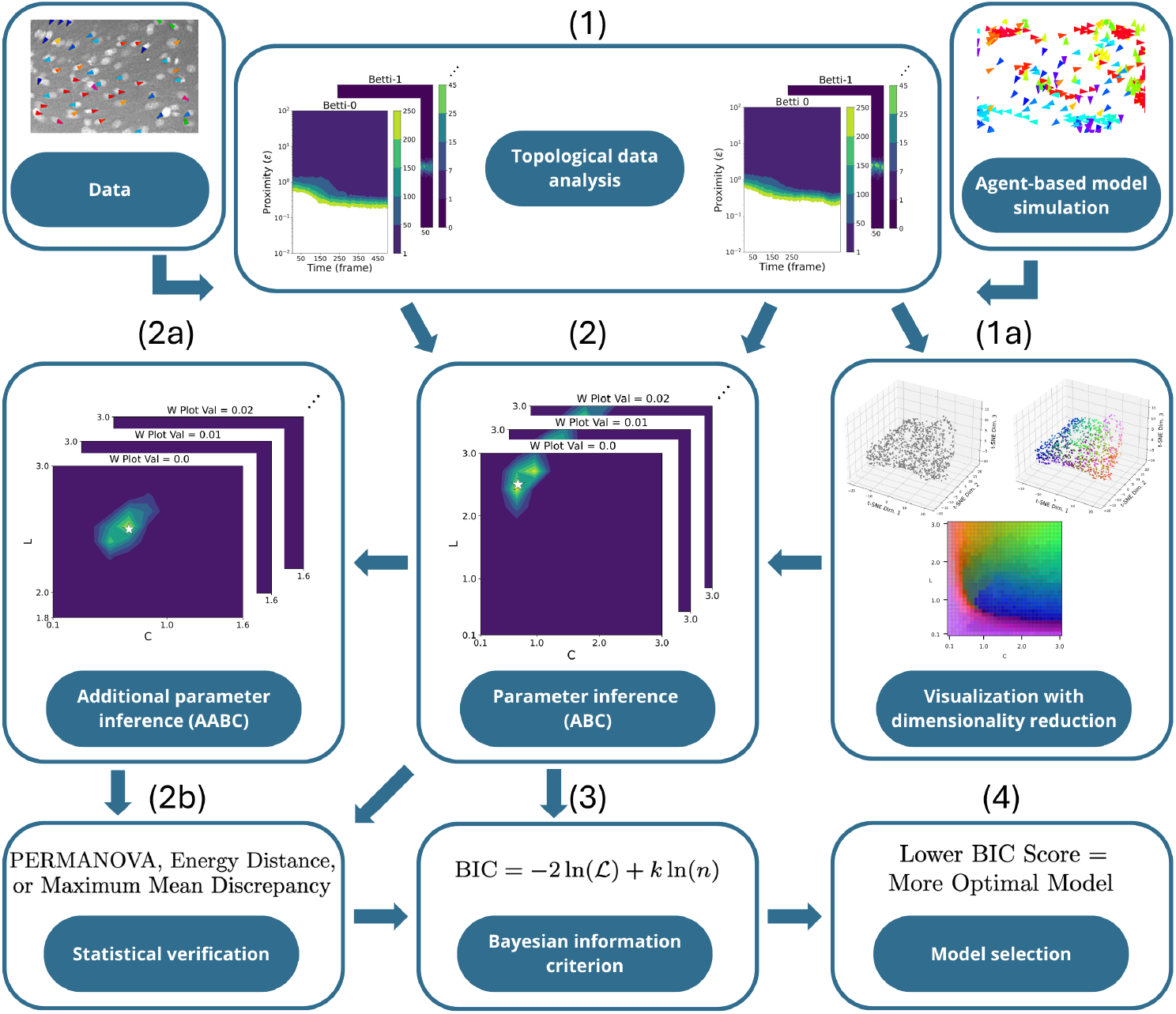
A computational pipeline for agent-based model selection. Our pipeline called TOPAZ has the following steps: (1) From data or an ABM simulation, topological data analysis in the form of Crocker plots. (1a) (Optional) Visualization with dimensionality reduction techniques such as t-SNE or principal component analysis. (2) Parameter inference techniques such as ABC. (2a) (Optional) Generation of additional samples using AABC for parameter inference if needed. (2b) (Optional) Assessment of distributional differences in posterior predictive summary statistics across models using PERMANOVA, Energy Distance, and/or MMD. (3) Bayesian information criterion score calculation. (4) Model selection based on lowest BIC score.

### Dimensionality reduction for visualization

To aid in addressing the inverse problem’s identifiability, t-SNE is employed as a nonlinear dimensional reduction method, which is particularly effective when computationally feasible. By reducing high-dimensional data to a three-dimensional plot, t-SNE helps visualize complex relationships more clearly. We simulated the new Alignment model for 30 equally spaced values of C and L ranging from 0.1 to 3.0, and 11 equally spaced values of W ranging from 0.0 to 0.1, resulting in 9,900 total combinations. For every simulation, we computed a Crocker matrix with size 100 time points×200 proximity parameters ×2 Betti numbers (zero and one). We applied t-SNE to reduce the 9,900 Crocker plots (example shown in (a) of Fig 2) with size 100 × 200 ×2 matrix to a 3 × 1 vector. A random 10% subset of the 9,900 points is visible in the t-SNE reduced space in (b) of Fig 2, with each point colored according to its 3D coordinates, converted into an RGB value (shown in (c) of Fig 2). This color data was then extracted to create eleven 30 ×30 2D t-SNE plots, one for each W value (examples shown in (d) and (e) of Fig 2; all eleven plots can be found in Supplementary Material Figure S2). For more visualization using t-SNE, see Supplementary Material Appendix S4 and Supplementary Material Figure S1.

**Fig 2.**
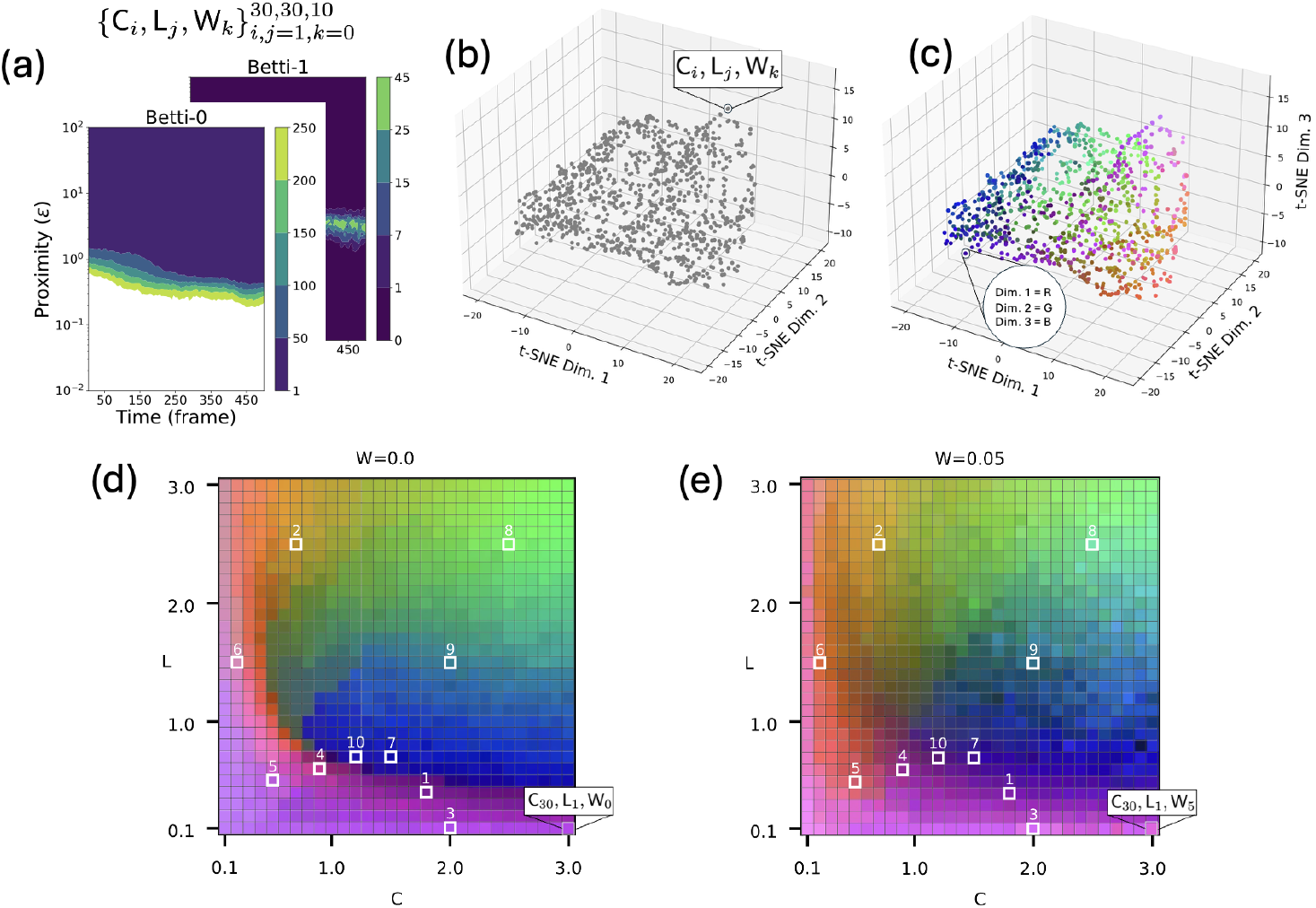
Visualization with dimensionality reduction using t-SNE. (a) Crocker plots of the ABM simulation. (b) Size 100× 200× 2 matrix reduced to a 3 ×1 vector. Only 10% of the 9,900 simulations are shown. (c) Each point is colored based on their 3D position where (Dim. 1, Dim. 2, Dim. 3) = (R, G, B). (d and e) A 2D extraction from the 3D plot. Each point in the 3D plot has a C, L, and W value. These are plotted in 2D grid of C and L for each value of W. Left (d) is the W=0 slice and right (e) is the W=0.05 slice. The ten selection points were chosen for further analysis to cover a diverse range of behaviors (see Table 1).

**Table 1.**
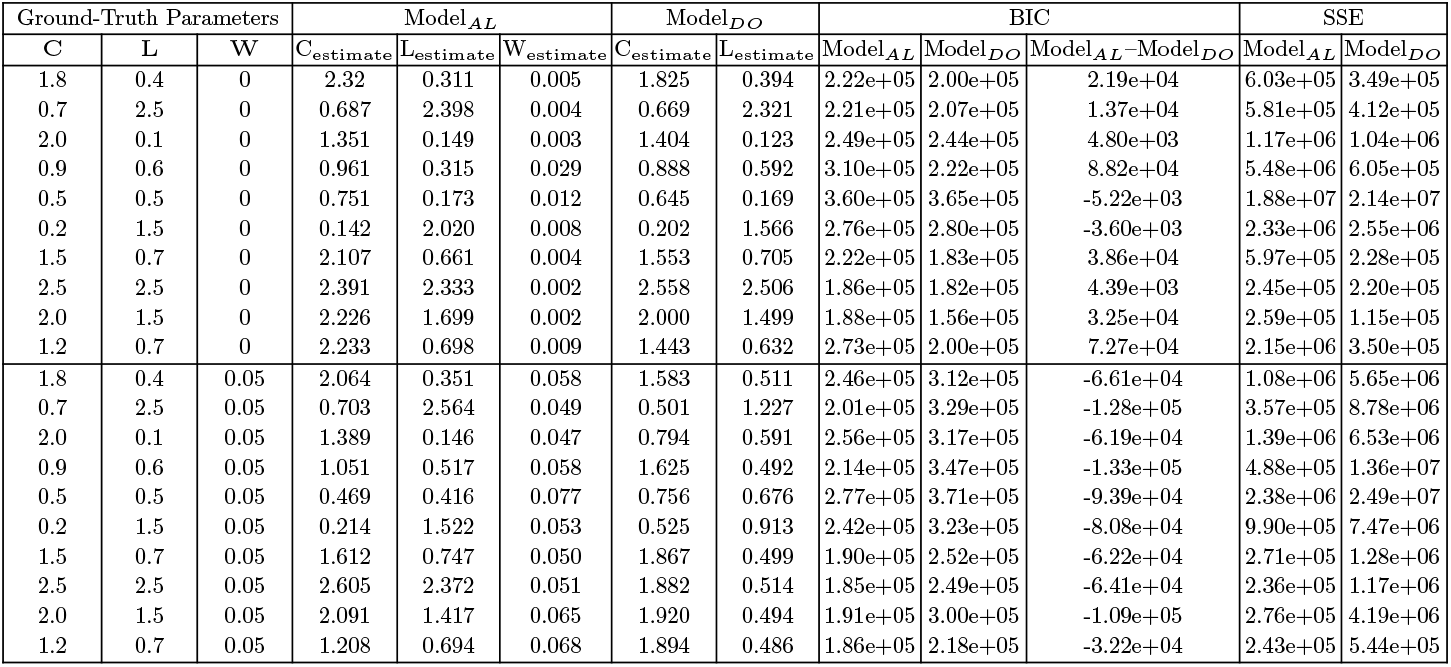
Model selection comparison results of Model_AL_ to Model_DO_. Results are summarized using AABC-estimated parameter values for both models, the BIC score and sum of squared errors (*SSE* ) for each model, and the difference of BIC scores. This is done for the ten ground truth samples chosen in (d) and (e) of Figure 2, corresponding to the selection of C, L, and W values in the first three columns.

## Results

### Simulation Study Design and Hypotheses

We developed a computational pipeline TOPAZ for selecting which ABM is the optimal fit for topological attributes derived from spatiotemporal cellular data. We tested this pipeline in a simulation study comparing fits of the ABM Model_AL_ to data simulated by the ABM Model_DO_, and also fits of the ABM Model_DO_ to data simulated by the ABM Model_AL_. Our hypothesis is that by analyzing the outputs of our TOPAZ framework, including t-SNE, *SSE*, BIC, nonparametric statistical tests (PERMANOVA, Energy Distance, and MMD), and posterior density plots, the user will be able to conclude when Model_AL_ can be selected as the better model. By leveraging BIC in addition to TDA, our TOPAZ pipeline penalizes for models with additional parameters and therefore only selects more complex ABMs when that complexity is represented in the topology of the data. Similarly, we also hypothesized that when data are simulated from the Model_DO_ model, TOPAZ will select Model_DO_ instead of Model_AL_ as the optimal model because the additional parameter in Model_AL_ does not provide a significantly better fit to the topological information in the Model_DO_ simulation, as determined by BIC (see step 4, Fig 1).

### Parameter Selection and Synthetic Data Generation

We selected a set of 10 ground-truth parameter vectors for each ABM corresponding to different values of (C, L, W) for Model_DO_ and Model_AL_, where Model_DO_ corresponds to W = 0 (Table 1, columns 1-3). Our ground-truth parameter choices were based on previous study by Nguyen et al. showing a wide representation of topologically distinct spatiotemporal features in simulations of the Model_DO_ model [12]. The same values of C and L were used for both Model_DO_ and Model_AL_ to isolate the effects of the added alignment framework. If we randomized all C, L, and W values, it would be harder to compare the effects of the added alignment parameter. Thus, by keeping C and L the same between models and only changing W, we can see the effect that our new alignment parameter has on the system. We chose W = 0.05 for simulating data for Model_AL_ (see parameter choices 1-10 in (d) and (e) of Fig 2). Representative snapshots of ABM simulations from Model_DO_ and Model_AL_ are shown for parameter choices 1 and 5 in Fig 3, visually exemplifying topological differences between the cases when W = 0 (No Alignment) and W = 0.05 (Alignment). See Supplementary Video S1-Supplementary Video S4 for the corresponding ABM simulation movies.

**Fig 3.**
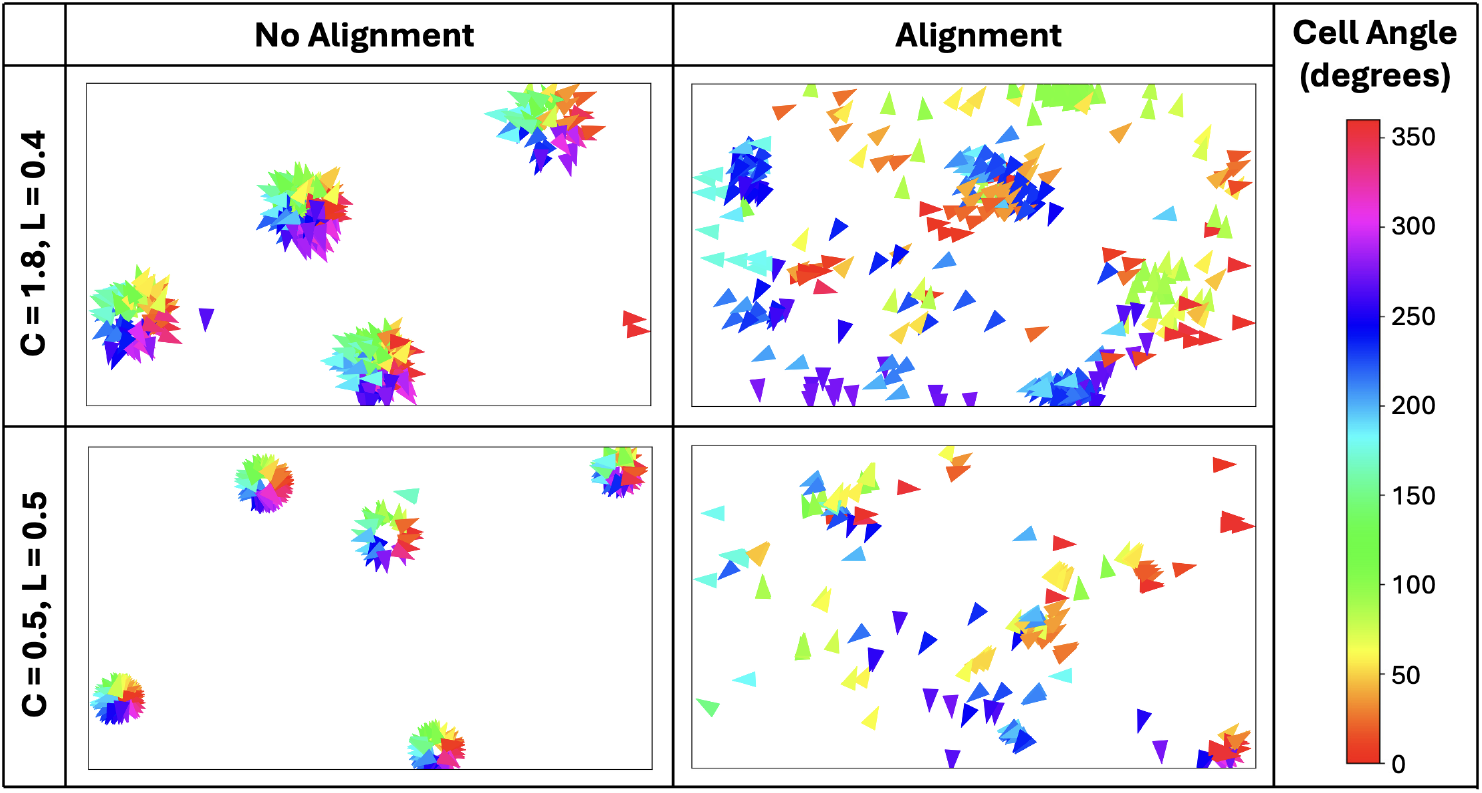
Visualization of Model_AL_ output for two different C and L combinations. Simulation snapshots are shown without alignment case (W=0.0, left) and with alignment case (W=0.05, right). The arrows and colors represent the direction the cells are moving. These snapshots are at the end of the simulation (t=120, last frame).

### Parameter Inference Using ABC and AABC

Using simulations generated from Model_DO_ or Model_AL_, we computed TDA summaries from the generated point clouds and then used them for parameter inference within ABC and AABC (steps 1, 2, and 2a in Fig 1). Visualizing the AABC posterior distributions exemplifies that the median of the posterior was close to the ground-truth parameter value when the ground-truth case was for W = 0 or W = 0.05 (Fig 4). The complete set of posterior plots for all W values is displayed in Supplementary Material Figure S3. Concordantly, the Crocker plots at the ground-truth and estimated parameter values (median of posterior) exemplify that the corresponding topological diagrams are visually similar (Fig 5).

**Fig 4.**
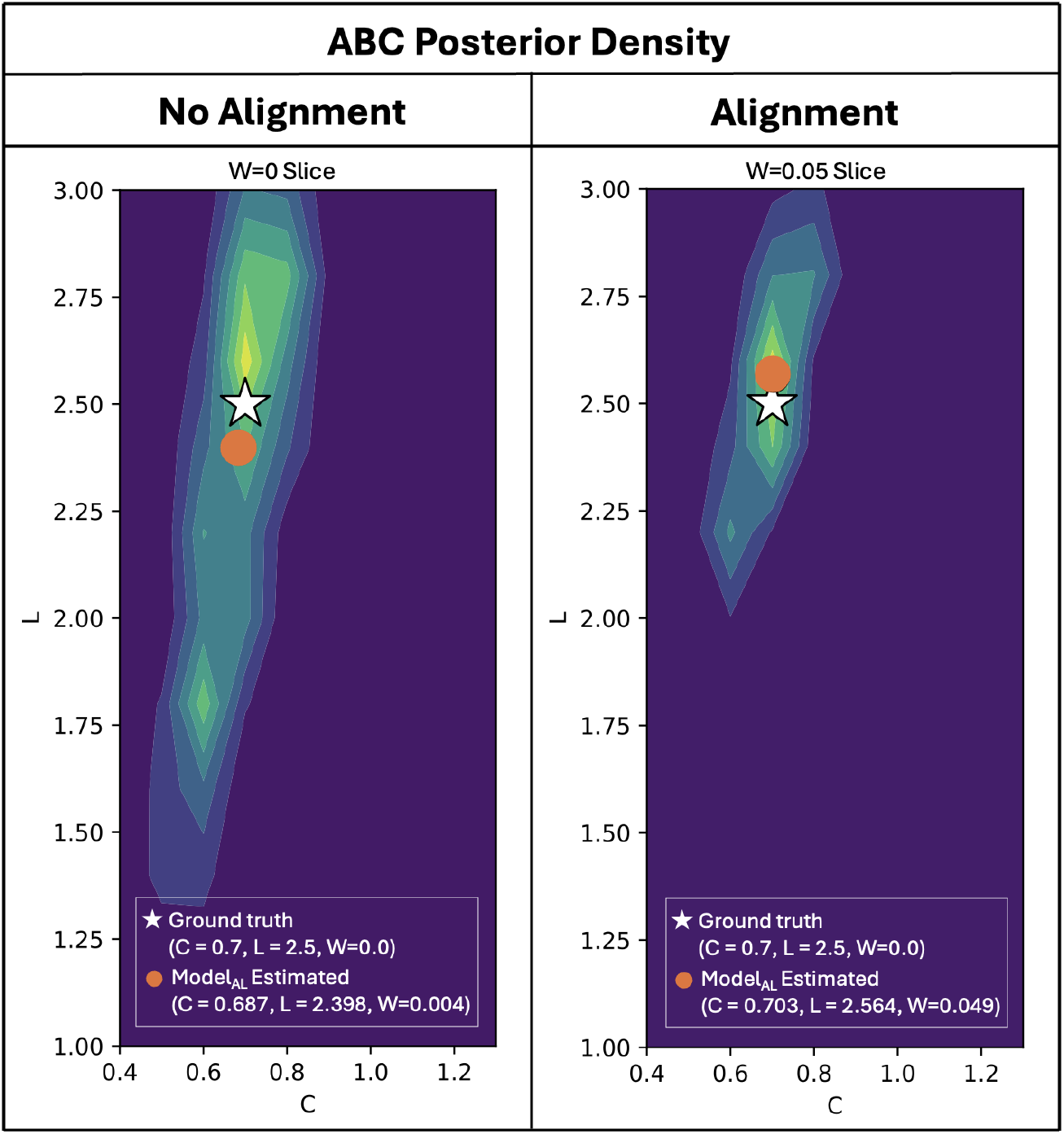
Examples of AABC posterior density plots for Model_AL_. The ground truth (white stars) and sample median AABC-estimated parameter values for Model_AL_ (orange dots) are displayed as well as the AABC posteriors for Model_AL_. Left and right use data generated with no alignment and alignment, respectively. For the sample median AABC-estimated parameter values for Model_DO_, see Table 1. The complete selection of AABC posterior density plots for Model_AL_ can be found in Supplementary Material Figure S3.

**Fig 5.**
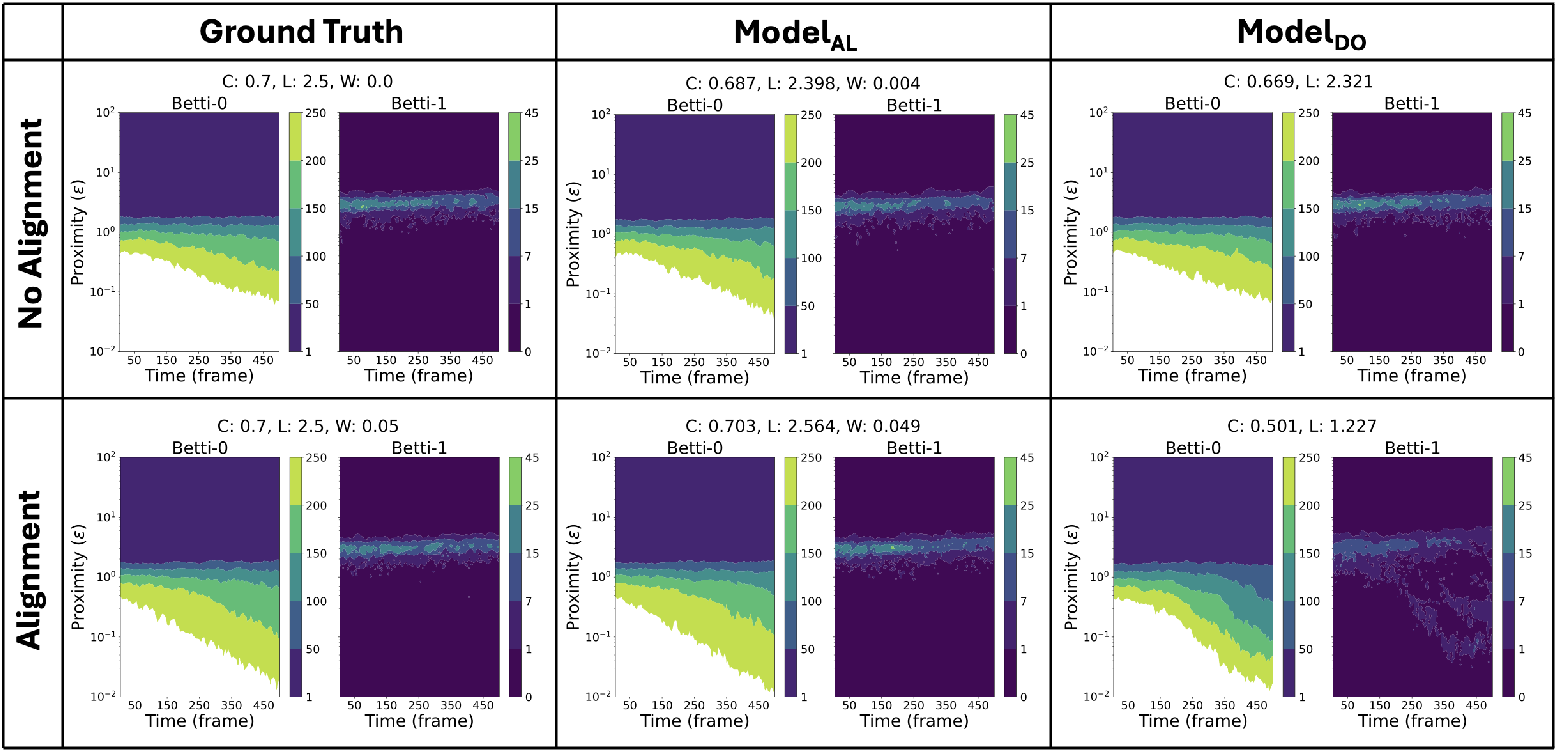
Crocker plots summarizing the topology of the ground truth and AABC simulations. For the ground truth values and AABC-estimated values for Model_AL_ and Model_DO_, we constructed Crocker plots for Betti-0 and Betti-1 (left, middle, and right, respectively) for the case where there is no alignment (W=0.0, top) and alignment (W=0.05, bottom). The color legend represents the contour levels corresponding to the number of simplices of degree 0 and 1 at each proximity value and frame number; counts exceeding 250 are shown in white for visualization purposes.

### Model Selection via BIC

To quantitatively and comprehensively summarize the visual observations we made, BIC and AIC scores were calculated for Model_AL_ and Model_DO_ against simulated data generated at each of the 10 chosen parameter vectors. Table 1 displays BIC scores as well as the difference between the Model_AL_ and Model_DO_ BIC scores for comparison (right-most column). The AIC results can be found in Supplementary Material Table S2. We found that the BIC for Model_AL_ was lowest for all the Model_AL_ simulated data and the BIC for Model_DO_ was lowest for all the Model_DO_ simulated data except two cases (see Supplementary Material Appendix S6). In these two cases, Model_AL_ identified alternative parameter combinations with small inferred values of *W* that yielded lower BIC values but did not more closely recover the ground-truth parameters, highlighting the difficulty of distinguishing between the two models when the inferred alignment strength is small. Overall, these results demonstrate that our TOPAZ pipeline can quantitatively compare ABMs and provide a framework for model selection for spatiotemporal cellular data.

## Discussion

We introduced TOPAZ, a novel methodology to facilitate model selection for ABMs. This algorithm integrates concepts from TDA, ABC, AABC, and statistical modeling to achieve this. We apply the Vietoris-Rips filtration to summarize the topological signature of an ABM simulation using agents’ output data. For a given dataset and library of candidate models, we use ABC and AABC to infer the posterior distribution of each considered model’s parameter values. The BIC then identifies which model best describes the data with a reasonable amount of complexity. Model selection facilitates the identification of relevant biological mechanisms in the presence of noisy data [25]. For our use of BIC, we use the sample median because it is a comparison to previous work [12], but we want to acknowledge that the sample mean may also be used and that the BIC is based on the Maximum Likelihood Estimation fundamentally. Although BIC is well aligned with the model-identification objective of the present study, alternative information criteria may be more appropriate in applications prioritizing predictive performance or involving different model structures and data regimes. In calculating BIC and AIC, we used a Gaussian error model on the Crocker-summary residuals, with the residual variance estimated as σ^2^ = *SSE/n*. This choice provides a scale-adaptive likelihood approximation for the present proof-of-concept study, but it does not address possible non-constant variance, correlation, or model-discrepancy structure in experimental single-cell data. Future applications of TOPAZ to experimental data could replace this homoscedastic Gaussian approximation with weighted or generalized least-squares error models, or with a fully Bayesian likelihood that includes an observation covariance or variance model, following established inverse-problem and uncertainty-quantification approaches [26–28].

We developed an extension of the D’Orsogna Model that includes an agent alignment force. The goal was to incorporate a new biophysical mechanism that is biologically grounded to create a more realistic model. This new model incorporates an alignment term that promotes directional coordination among neighboring cells. We found that simulations from the new model are more likely to exhibit fluidization behavior, i.e., the formation of groups of cells moving in coherent directions parallel or antiparallel to adjacent groups. This recapitulates experimental observations of confluent fibroblast populations, where cells self-organize into dynamic, parallel/anti-parallel streams [12]. We then considered a suite of ABM simulations that were simulated from different parameter values. We showed that the TOPAZ algorithm can be used as a quantitative tool for model selection and parameter estimation utilizing topological summaries of spatiotemporal single-cell data. It could also be applied to other ABMs with spatiotemporal data in other areas such as ecology or epidemiology. This helped answer the question of can we use quantitative or topological metrics to tell if our extended model is actually a better model.

While we applied this flexible framework to a model of biological swarming, it is broadly applicable for many modeling approaches in molecular and cellular biology [29]. We designed our computational pipeline with future applications in mind, particularly spatial single-cell datasets that incorporate transcriptomic and proteomic measurements. In particular, intracellular signaling pathways regulate dynamics of the actin cytoskeleton, thereby breaking symmetry of intracellular forces to drive both individual- and population-level cell behavior [30–32]. For *in vivo* settings, dynamic changes to the tissue microenvironment might also need to be considered [33–35]. In these scenarios, our model selection framework can be readily applied by incorporating microenvironment variables into an ABM, similar to other recently developed multiscale models [36, 37]. Since the TOPAZ framework operates on arbitrary multidimensional simulation outputs, it can be used to select among models that track both spatial trajectories and intracellular signaling states [38]. Such an analysis could reveal the relationship between biochemical pathway activation and collective cellular migration [39, 40].

We incorporated many simple algorithms into our first pass of the TOPAZ algorithm to demonstrate its utility. For example, we summarized the time-varying topological signature of ABM simulation using Crocker plots and performed the rejection algorithm for our ABC and AABC computations. We plan to implement other methods in future work to enhance the TOPAZ algorithm. Vineyards and the Crocker stack have been introduced as stable approaches to topologically represent time-varying data [41, 42]. We can also use the principal component analysis or t-distributed stochastic neighbor embedding dimensionality results in place of our full Crocker plots as a new summary statistic instead of only for visualization purposes. However, this is beyond the scope of this paper and will be investigated in future studies. For ABC and AABC parameter inference, we successfully used the ABC and AABC algorithms to estimate 3 parameter values. For future work, we will investigate if these approaches may be able to estimate more parameter values and if the error in the parameter estimation is due to the AABC algorithm, the coarseness of the grid, or other factors. In this scenario, we may turn to more data-efficient algorithms, including Sequential Monte Carlo methods [43]. Since the TOPAZ method has so far been validated only on simulated data, an important direction for future work is assessing its robustness in the presence of experimental noise, as demonstrated in related workflows such as Nguyen et al. [12].

Although many alternative model families could have been considered for model selection, in this initial study we focused on the D’Orsogna model used in Nguyen et al. [12] and an intentionally extended model that only adds one biological mechanism - the additional alignment term. Because the baseline D’Orsogna model is already known to reproduce key qualitative features of our experimental system, we viewed the alignment extension as a biologically motivated next step toward improving its ability to capture experimental observations. Our current work establishes this extension and evaluates the TOPAZ framework under controlled conditions using simulated data.

Importantly, comparing two closely related models provides a stringent test of the model-selection component of TOPAZ. Distinguishing between models that differ only subtly in their mechanistic structure is inherently challenging; demonstrating that the framework can resolve these small differences increases confidence in its ability to discriminate among more disparate or non-nested models. In future work we will explore the application of the TOPAZ framework to a broader set of candidate models, including further extensions of the D’Orsogna family as well as models that incorporate qualitatively different interaction rules.

Topological data analysis, and persistent homology in particular, provides robust, multiscale summaries that capture spatial organization, gradients, and connectivity. The ABMs we investigate here demonstrate that such topological summaries offer an effective interface between complex single-cell data and mechanistic models across biological scales. While the current study is limited to cell tracking data and a minimal ABM, TOPAZ is an extensible model selection tool for ABMs that integrates TDA, thereby enabling the extraction of mechanistic insight from the type of noisy high-dimensional measurements that are characteristic of spatiotemporal single-cell datasets.

## Methods

### Topological data analysis

Homology is a fundamental concept in algebraic topology, quantifying the topological structure of data by identifying the number of *n*-dimensional holes within a given space. Persistent homology (PH) extends this notion by capturing topological features across multiple scales, providing a multi-scale summary of the shape of data. In particular, PH can be used to quantify changes in the topological structure of biological data, such as the collective motion of cells over time, by calculating Betti numbers. For a comprehensive introduction to PH and its application to biological systems, refer to [12].

To investigate the topological structure of cell motion, we compute the persistent homology of point clouds representing the time-varying locations and orientations of a population of cells. Each frame of the simulation is represented as a point cloud, denoted 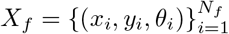, where *N*_*f*_ is the number of cells in frame *f* . In this context, each point represents a 0-simplex, with coordinates *x*_*i*_, *y*_*i*_, and *θ*_*i*_ corresponding to the spatial location and orientation of the *i*-th cell at time *f* .

To analyze the topology of these point clouds, we use the Vietoris-Rips filtration, which generates simplicial complexes from point clouds by progressively adding higher-dimensional simplices (edges, triangles, etc.) based on a proximity parameter *ε*. The Vietoris-Rips complex 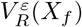 is constructed by connecting any set of *k* + 1 points within distance *ε* of each other. The distance between two points is computed as the Euclidean distance in the *x* and *y* dimensions, plus the angular distance in the *θ* dimension, where angular distances are calculated as the shortest path on a circle, i.e., *θ*_1_ *− θ*_2_ = min(*θ*_1_, *θ*_2_) *−* max(*θ*_1_, *θ*_2_) + 360.

By taking an increasing sequence of proximity parameters *T* = (*ε*_0_, *ε*_1_, …, *ε*_*m*_), we generate a nested family of simplicial complexes, known as a filtration. For each *ε*, we compute the *k*-th Betti number of the associated complex 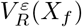, which counts the number of *k*-dimensional holes (or topological features) in the complex. Specifically, the *k*-th Betti number is defined as the rank of the *k*-th homology group, which quantifies the number of *k*-dimensional holes enclosed by 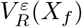 [44].

To summarize the persistent homology of dynamic point clouds, we compute Betti curves for each time step and then concatenate them into a single matrix, which summarizes the homology of the entire time series. This matrix is known as a *Crocker plot* [45]. Crocker plots are a powerful tool for visualizing the evolution of Betti numbers over time by plotting the level sets or contours of concatenated Betti curves. These plots have been used to study the topology of biological systems, such as cell trajectories, by encoding topological changes at multiple spatial scales [12].

For all persistent homology computations, we used the Ripser package, a highly optimized C++ library for computing persistent homology [46]. Ripser provides state-of-the-art performance by reducing both memory consumption and computation time. We used the Python port of Ripser, available through the Scikit-TDA library, to implement all PH calculations in this analysis [47].

### Approximate Bayesian computation

In contrast to frequentist methods, which provide point estimates for model parameters, Bayesian inference estimates a probability distribution over parameters conditioned on observed data. Specifically, the goal is to estimate the posterior distribution *p*(***q***|**Y**^*o*^), where ***q*** is the vector of parameters and **Y**^*o*^ represents the observed data. According to Bayes’ theorem, this posterior is given by:

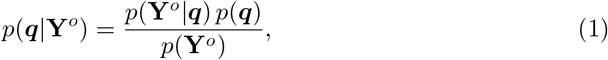

where *p*(**Y**^*o*^|***q***) is the likelihood function, *p*(***q***) is the prior distribution, and *p*(**Y**^*o*^) is the marginal likelihood:

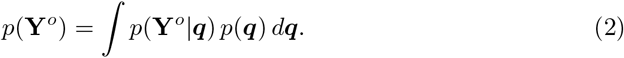

While the marginal likelihood plays a crucial role in model comparison, it is often computationally intractable due to the high-dimensional integration involved. Fortunately, since it does not depend on the parameters ***q***, it can be treated as a constant during parameter estimation and safely omitted when comparing posterior densities across parameter values. However, computing the marginal likelihood is still necessary for formal model selection tasks.

A major challenge in Bayesian inference arises when the likelihood function *p*(**Y**^*o*^|***q***) cannot be evaluated directly. This situation is common in complex or stochastic models [15, 48, 49]. In such cases, ABC provides a powerful alternative. ABC was originally introduced in Tavar*é* et al. as a likelihood-free, rejection-based method [50]. Since then, ABC has been extended and refined in various studies [48, 51, 52].

Comprehensive overviews of ABC methods are available in [53, 54]. In this study, we use the standard rejection-based ABC algorithm, also known as the prototype rejection-ABC algorithm [53], which we refer to simply as the ABC algorithm.

ABC approximates the likelihood by generating simulated datasets **Ŷ** from the model using parameters drawn from the prior distribution. These simulated datasets are then compared to the observed data **Y**^*o*^ using a distance function *ρ*(**Y**^*o*^, **Ŷ** ). Parameter samples for which the simulated output is sufficiently close to the observed data (i.e., within a specified tolerance *τ* ) are accepted. As described in [12], the posterior can be approximated as:

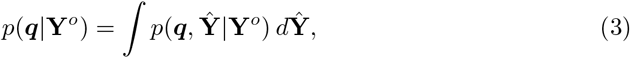

with the joint distribution given by:

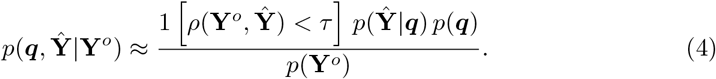

In our approach, we use Crocker matrices of Betti numbers 0 and 1 as summary statistics to characterize model behavior, as done in previous work [12]. Specifically, the observed data **Y**^*o*^ is replaced by the ground-truth Crocker matrix 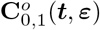, and the simulated data **Ŷ** is represented by **Ĉ** _0,1_(***t, ε***; ***q***), which is generated using candidate parameter values. To quantify similarity, we employ the sum of absolute errors as our distance metric. A sample ***q*** is accepted if it satisfies:

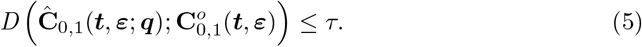

where

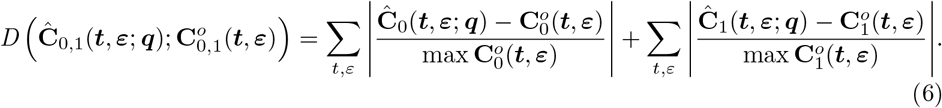

We assume that only the upper and lower bounds of the parameters C and L are known, and we use a joint uniform prior over the interval [0.1, 3.0]. A total of *N*_*s*_ = 10,000 samples are drawn from this prior all with the same fixed initial conditions.

Choosing the tolerance *τ* is crucial for balancing accuracy and acceptance rate. However, in this work, we choose to accept only the top 1% samples with best error values. The accepted samples form the so-called ABC-posterior density. To compare the ground-truth and the estimated, we compute the median for each parameter from the accepted samples.

### Approximate approximate Bayesian computation

As an extension of ABC, we employ AABC to efficiently generate additional samples for parameter inference. AABC follows the same initial and final steps as ABC, beginning with a finite set of forward simulations and concluding with a rejection-based approximation to the posterior distribution. Its primary advantage lies in its ability to produce a substantially larger number of approximate samples by leveraging an initial set of simulated data, thereby reducing the computational burden associated with repeated forward simulations [55].

We first performed ABC using *N*_*s*_ = 10,000 simulations to obtain posterior density estimates for each model. To concentrate subsequent sampling in regions of non-negligible posterior support, we restricted the original 30×30×11 parameter grid to a smaller grid containing all parameter combinations for which the posterior density was nonzero for either Model_DO_ or Model_AL_ in each row of Table 1. This restriction allows computational effort to be focused on regions of interest rather than distributed uniformly across the full parameter space.

Following this grid refinement, we applied the first stage of the AABC algorithm by generating an expanded set of original simulations within the reduced parameter space. The total number of simulations was scaled proportionally to the size of the reduced grid and then multiplied by a factor of ten relative to the original ABC sample size; for example, if the restricted grid occupied half of the original parameter space, we generated 50,000 simulations. These simulations served as the basis for constructing additional approximate samples without further evaluations of the underlying agent-based model.

New samples were generated using the AABC procedure described in Buzbas et al. [55]. For each parameter proposal, the *k* = 5 nearest neighbors were identified using Euclidean distance in parameter space, where each Crocker plot was treated as a single data point. Parameter values were then averaged using Epanechnikov kernel weights to produce approximate posterior samples. This process was repeated iteratively, with parameter estimates, error metrics, and BIC values recomputed after each batch of newly generated samples. Convergence was assessed by comparing the cumulative approximate posterior distributions at successive sampling checkpoints. For each parameter, changes in five posterior quantiles (0.05, 0.25, 0.50, 0.75, and 0.95) and the first Wasserstein distance between successive marginal posterior distributions were calculated and normalized by the corresponding prior parameter range. A checkpoint comparison passed the convergence criterion when both the maximum normalized quantile change and the maximum normalized Wasserstein distance across all parameters were less than 0.01. Convergence was declared after this criterion was satisfied for three consecutive checkpoint comparisons. Convergence behavior is illustrated in Supplementary Material Figure S4.

### Bayesian information criterion

BIC, also known as the Schwarz information criterion, is a widely used criterion for model selection and comparison, particularly in the context of statistical modeling and machine learning. It is employed to assess the trade-off between model complexity and goodness of fit, providing a way to avoid overfitting by penalizing the inclusion of additional parameters in the model.

The BIC is defined as:

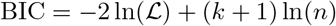

where ℒ is the likelihood of the model given the data, *k* is the number of parameters in the model, and *n* is the number of data points. The ln(ℒ) is defined as:

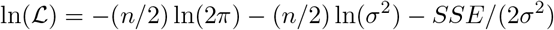

where *σ*^2^ = *SSE/n*.

The penalty term (*k* + 1) ln(*n*) discourages overly complex models by penalizing models with more parameters. The ABC rejection procedure and BIC computation are summarized in Algorithm 1 and the AABC rejection procedure and BIC computation are summarized in Algorithm 2.

#### Algorithm 1 Approximate Bayesian computation rejection and Bayesian information criterion algorithm

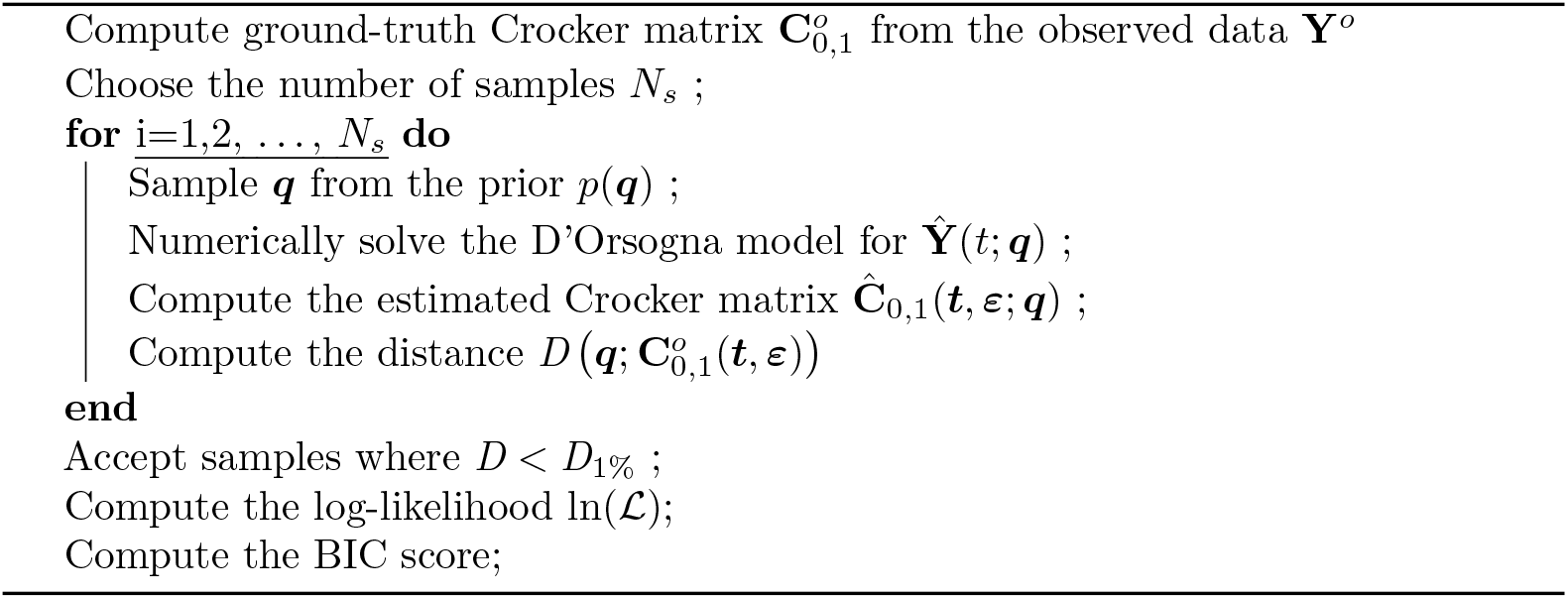

#### Algorithm 2 Approximate approximate Bayesian computation rejection and Bayesian information criterion algorithm

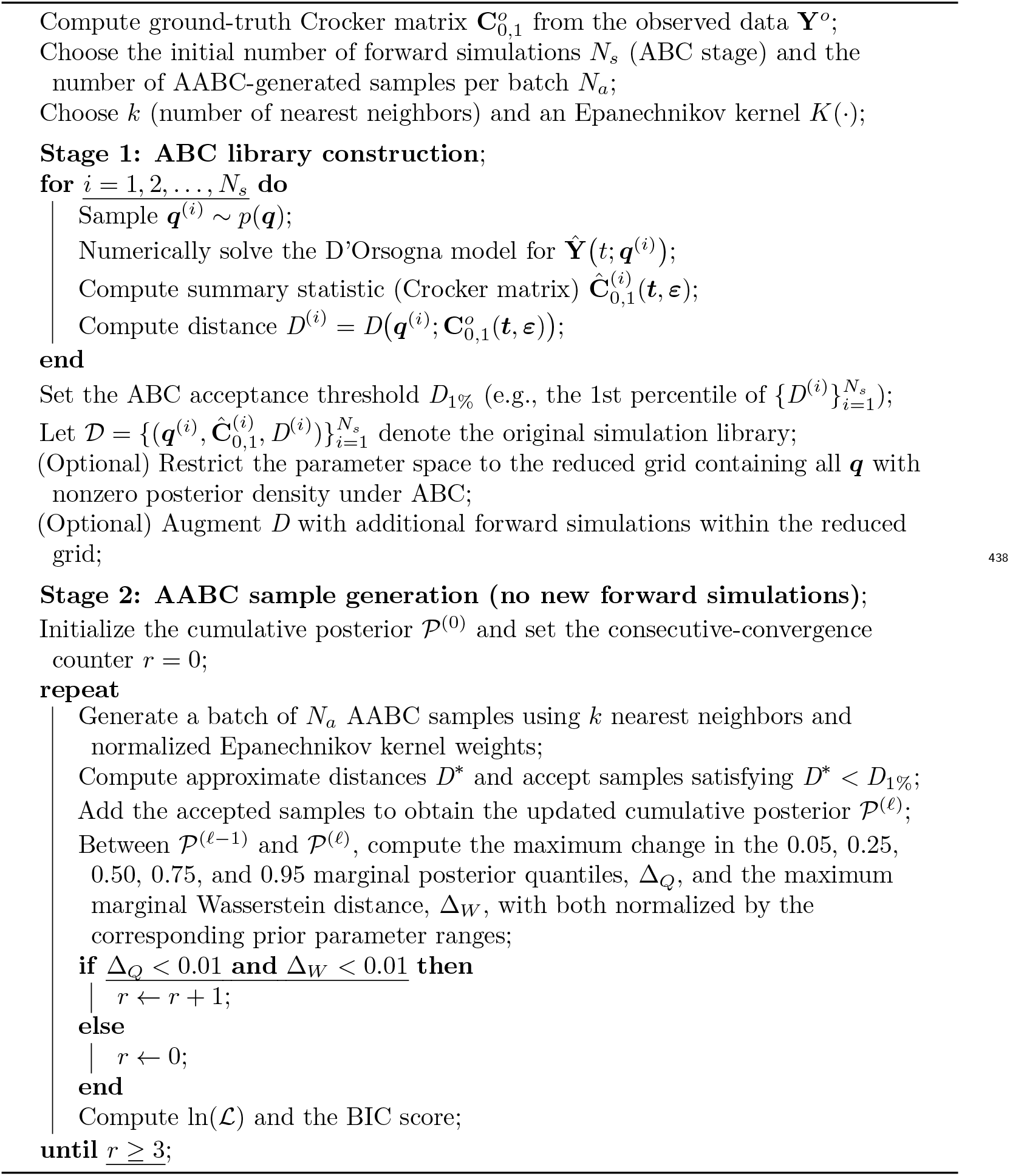

## Supporting information

**Supplementary Material Appendix S1. D’Orsogna and Alignment models. Supplementary Material Appendix S2. Experimental set-up**.

**Supplementary Material Appendix S3. Statistical evaluation for model comparison**.

**Supplementary Material Appendix S4. Nearest neighbor t-SNE visualizations**.

**Supplementary Material Appendix S5. Akaike information criterion**.

**Supplementary Material Appendix S6. Case Analysis: When Model**_**AL**_ **outperforms Model**_**DO**_.

**Supplementary Material Table S1. Model**_**AL**_ **versus Model**_**DO**_ **Crocker plot distribution statistical difference results**.

**Supplementary Material Table S2. Model**_**AL**_ **versus Model**_**DO**_ **Model selection comparison results of Model**_**AL**_ **to Model**_**DO**_ **using AIC**.

**Supplementary Material Figure S1. T-SNE nearest-neighbor visualizations**. T-SNE nearest-neighbor visualizations for the selected (C,L) pairs from Table 1 when W=0.0 and W=0.05. All points where W=0 are in red and similarly all points where W=0.05 are in blue. Each plot has one selected (C,L) pair at W=0.0 and its nearest neighbor where W=0.05 for the left two columns and the reverse on the right.

**Supplementary Material Figure S2. Full selection of 2d t-sne extractions**. The full selection of the Betti-0 and Betti-1 t-SNE colormaps, showcasing an example of no alignment (W=0.0, top left) and different levels of alignment (W=0.01-W=0.1). The colors for each (C, L, W) combination are the colors that were generated in the 3D SNE plot ((c) of Figure 2) for the corresponding (C, L, W) point.

**Supplementary Material Figure S3. Full selection of AABC posterior density plots for Model**_**AL**_. Top and bottom use data generated with no alignment (W=0.0) and alignment (W=0.05), respectively. The white star represents the ground-truth values whereas the orange dots represent the sample median AABC-estimated values for Model_AL_.

**Supplementary Material Figure S4. AABC Convergence plots**. The full selection of convergence plots for each row of Table 1 for both Model_DO_ and Model_AL_ when the ground truth value for W is 0 and 0.05 for a total of four convergence plots per row. The AABC method was used repeatedly to generate more samples until convergence has been met for each row of Table 1.

**Supplementary Video S1. Full simulation gif with (C, L, W)=(1.8, 0.4, 0.0)**. Video simulation from t=20-120. The arrows and colors represent the direction the cells are moving. Since W=0, this represents a case without alignment.

**Supplementary Video S2. Full simulation gif with (C, L, W)=(1.8, 0.4, 0.05)**. Video simulation from t=20-120. The arrows and colors represent the direction the cells are moving. Since W=0.05, this represents a case with alignment.

**Supplementary Video S3. Full simulation gif with (C, L, W)=(0.5, 0.5, 0.0)**. Video simulation from t=20-120. The arrows and colors represent the direction the cells are moving. Since W=0, this represents a case without alignment.

**Supplementary Video S4. Full simulation gif with (C, L, W)=(0.5, 0.5, 0.05)**. Video simulation from t=20-120. The arrows and colors represent the direction the cells are moving. Since W=0.05, this represents a case with alignment.

## Data and Code Availability

All code used to implement the TOPAZ framework, including simulation scripts, topological data analysis modules, and example notebooks, is publicly available at: https://github.com/alyswenz31/TOPAZ

