## Supplementary Material for "Topologically-based parameter inference for agent-based model selection from spatiotemporal cellular data"

### Supplementary Material Appendix 1. D'Orsogna and Alignment models.

#### D'Orsogna model

The D'Orsogna model originally published in [1] is a mathematical framework used to describe the dynamics of self-propelled particles, such as those found in biological systems like swarming or flocking behaviors. It involves parameters that govern attraction and repulsion among agents. The system of ordinary differential equations modeling the D'Orsogna swarm behavior for $1 \leq i, j \leq N$ agents are detailed in (1).

$$\begin{aligned} \dot{\mathbf{x}}_i &= \mathbf{v} \\ m\dot{\mathbf{v}}_i &= \left(\alpha - \beta\|\mathbf{v}_i\|^2\right)\mathbf{v}_i - \nabla_{\mathbf{x}_i} U_i \\ U_i &= \sum_{j\neq i}^{N} C_r e^{-\|\mathbf{x}_j - \mathbf{x}_i\|/l_r} - C_a e^{-\|\mathbf{x}_j - \mathbf{x}_i\|/l_a} \end{aligned} \tag{1}$$

In the D'Orsogna model, $\alpha$ and $\beta$ represent the self-propulsion and friction for each agent, respectively. We have fixed $\alpha$=1 and $\beta = 0.5$ to be consistent with past work [1–3]. If no other forces are acting on an agent, the agent will asymptotically approach a constant speed of $\|\mathbf{v}_i\| = \sqrt{\alpha/\beta}$. In capturing the repulsion and attraction experienced by each cell, we use a Morse potential $U_i$ for the $i^{\text{th}}$ agent. The Morse potential, while originally used to describe the potential energy of a diatomic molecule, provides a unique advantage in that it details the energy of an anharmonic oscillator; i.e., agents very close to $\mathbf{x}_i$ will produce a strong repulsive effect, but far away cells will produce a weakening attractive effect. The parameters representing the repulsive and attractive magnitudes are $C_r$ and $C_a$, respectively, and the parameters accounting for the spatial sensitivity of repulsion and attraction are $l_r$ and $l_a$, respectively. Often the ratios $\mathrm{C} := C_r/C_a$ and $\mathrm{L} := l_r/l_a$ are used in the plots of this paper and the phase diagram of Morse potential in D'Orsogna's paper. In a system of two particles, the zero force steady state distance is

$$\|\mathbf{x}_j - \mathbf{x}_i\| = l_r \frac{\ln(\mathrm{C}/\mathrm{L})}{1-\mathrm{L}} = l_a \mathrm{L} \frac{\ln(\mathrm{C}/\mathrm{L})}{1-\mathrm{L}}.$$

For there to be a physically meaningful steady state distance between two particles, we require that $\mathrm{L} < 1 < \mathrm{C}$ or $\mathrm{C} < 1 < \mathrm{L}$. For $\mathrm{C} < 1 < \mathrm{L}$, the Morse potential exhibits a maximum and there is no stable equilibrium. D'Orsogna et al. further describe in [1] systems beyond the parameters of (C, L) that are *H-stable* and *catastrophic*. The long-term patterns of H-stable systems scale with the number of particles; catastrophic systems collapse as the number of particles increase to infinity. Furthermore, the H-stable regions for their model must satisfy the following conditions: (I) $\mathrm{C} > 1$ and (II) lie above the separatrix $\mathrm{CL}^2 = 1$; all other regions are catastrophic.

#### Alignment model

In (1), we define the force of motion for an agent as

$$\mathbf{F}_i := (\alpha - \beta\|\mathbf{v}_i\|^2)\mathbf{v}_i - \nabla_{x_i} U_i. \tag{2}$$

For our implementation, we seek to retain the primary forces which act on an agent; however, we want to extend the D'Orsogna model to include *alignment* $\mathbf{A}_i$ as an additional factor which changes velocity. We model alignment to fulfill the following assumptions:

- **Fluidization**: Neighboring agents will seek to move in parallel even if they are traveling in opposite directions. Perpendicular motion is the only instance where $\mathbf{A}_i = 0$.
- **Momentum**: Agents are given their own mass, $m_i$. Agents with greater momentum are more difficult to align; neighboring agents with greater momentum are more efficacious in affecting alignment.
- **Collective**: An agent's alignment is more greatly affected by the total alignment of all neighboring agents within a radius $R$.

We define the sensitivity to alignment as $\mathrm{W} \geq 0$. We define $0 \leq \Delta\theta_{in} \leq \pi$ as the angle between the agent vector $\mathbf{v}_i$ and its neighbor $\mathbf{v}_n$. Let $\chi_{2R}(\|\mathbf{x}_n - \mathbf{x}_i\|)$ be the characteristic function of a neighboring agent $\mathbf{x}_n$ being within a radial distance $R$ from the agent $\mathbf{x}_i$. Therefore, we can model our assumptions in the following way

$$\begin{aligned} \mathbf{A}_i &= \mathrm{W}\sum_{n=1}^{N} \chi_{2R}(\|\mathbf{x}_n - \mathbf{x}_i\|)\cos(\Delta\theta_{in})\mathbf{v}_n \\ &= \mathrm{W}\sum_{n=1}^{N} \chi_{2R}(\|\mathbf{x}_n - \mathbf{x}_i\|)\frac{\|\mathbf{v}_n\|}{\|\mathbf{v}_i\|}\mathrm{proj}_{\mathbf{v}_n}\mathbf{v}_i \end{aligned} \tag{3}$$

Accounting for the mass of each agent $m_i$, we include the 2D components of alignment and set up a polar representation of acceleration.

$$\begin{aligned} \begin{pmatrix} \dot{v}_{x,i} \\ \dot{v}_{y,i} \end{pmatrix} &= \frac{1}{m_i}\begin{pmatrix} F_{x,i} \\ F_{y,i} \end{pmatrix} + \begin{pmatrix} A_{x,i} \\ A_{y,i} \end{pmatrix} \\ &= \begin{pmatrix} \cos(\theta_i) & -\sin(\theta_i) \\ \sin(\theta_i) & \cos(\theta_i) \end{pmatrix}\begin{pmatrix} \dot{r}_i \\ r_i\dot{\theta}_i \end{pmatrix} \end{aligned} \tag{4}$$

When solving for $\dot{r}_i$ and $r_i\dot{\theta}_i$, it becomes clear that in the current formulation, alignment provides some change in the magnitude of velocity. Therefore, if we want to retain the assumption that the D'Orsogna forces are the only forces that change the magnitude of velocity of an agent, we need to scale alignment such that $\dot{r}_i = 0$. In order to project the alignment $\mathbf{A}_i$ onto the $r_i\dot{\theta}_i$-space, our scaled alignment contribution is now

$$\begin{aligned} \dot{\mathbf{v}}_i &= \frac{1}{m_i}\mathbf{F}_i + \mathbf{S}(\theta_i)\mathbf{A}_i \\ \mathbf{S}(\theta_i) &= \begin{pmatrix} \sin^2(\theta_i) & -\cos(\theta_i)\sin(\theta_i) \\ -\cos(\theta_i)\sin(\theta_i) & \cos^2(\theta_i) \end{pmatrix} \end{aligned} \tag{5}$$

### Supplementary Material Appendix 2. Experimental set-up.

For each model simulation, we used the same fixed initial condition for position and velocity for each cell. This was randomly generated from a uniform distribution. Each simulation had the same length of time ($t_0 = 1, t_f = 21, \Delta t = 1/6$) and the same number of agents ($\#_{agents} = 300$). When calculating the Crocker plots, we used the last 100 time points of the simulations. The total time was 120 time points and the first 10 time points were used as a burn-in and the second 10 time points were used to calculate the velocity of each cell.

### Supplementary Material Appendix 3. Statistical evaluation for model comparison.

#### Permutational Multivariate Analysis of Variance

Permutational Multivariate Analysis of Variance (PERMANOVA) is a nonparametric statistical test used to assess differences between groups based on multivariate data. Unlike traditional analysis of variance (ANOVA), which relies on assumptions of multivariate normality and homogeneity of variances, PERMANOVA operates directly on a distance or dissimilarity matrix, making it well suited for complex, high-dimensional, or non-Euclidean data.

PERMANOVA partitions the total variation in the distance matrix into within-group and between-group components and computes a pseudo-$F$ statistic analogous to the classical ANOVA $F$-statistic. Given a distance matrix $D$, the pseudo-$F$ statistic is defined as

$$F = \frac{SS_{\text{between}}/(g-1)}{SS_{\text{within}}/(n-g)},$$

where $SS_{\text{between}}$ and $SS_{\text{within}}$ are the sums of squares associated with between-group and within-group variation, respectively, $g$ is the number of groups, and $n$ is the total number of observations.

Statistical significance is assessed through a permutation procedure rather than reliance on a known parametric distribution. Group labels are randomly permuted a large number of times, and the pseudo-$F$ statistic is recomputed for each permutation. The resulting empirical distribution is used to estimate a $p$-value as

$$p = \frac{1 + \sum_{i=1}^{N} \mathbb{I}(F_i \geq F_{\text{obs}})}{1+N},$$

where $F_{\text{obs}}$ is the observed pseudo-$F$ statistic, $F_i$ denotes the statistic from the $i$-th permutation, $N$ is the total number of permutations, and $\mathbb{I}(\cdot)$ is the indicator function. For our test, we used $N$=999 permutations.

Because PERMANOVA is based on distances, its results depend on the choice of distance metric, and it is sensitive to differences in group dispersion. Nonetheless, it provides a flexible and robust framework for testing group-level differences in multivariate settings and is commonly used in ecological, biological, and other data-intensive applications.

#### Energy distance statistical test

The Energy Distance test is a nonparametric statistical test used to assess whether two samples are drawn from the same underlying probability distribution. It is based on the concept of statistical energy and provides a measure of dissimilarity between distributions that is sensitive to differences in location, scale, and overall distributional shape. The test is particularly well suited for multivariate data and does not rely on parametric assumptions such as normality.

Let $X = \{X_1, \ldots, X_n\}$ and $Y = \{Y_1, \ldots, Y_m\}$ denote two independent samples in $\mathbb{R}^d$. The population energy distance between the distributions of $X$ and $Y$ is defined as

$$\mathcal{E}(X,Y) = 2\,\mathbb{E}\|X-Y\| - \mathbb{E}\|X-X'\| - \mathbb{E}\|Y-Y'\|,$$

where $X'$ and $Y'$ are independent and identically distributed copies of $X$ and $Y$, respectively, and $\|\cdot\|$ denotes the Euclidean norm.

An unbiased sample-based estimator of the energy distance is given by

$$\hat{\mathcal{E}}_{n,m} = \frac{2}{nm} \sum_{i=1}^{n} \sum_{j=1}^{m} \|X_i - Y_j\| - \frac{1}{n(n-1)} \sum_{i \neq i'} \|X_i - X_{i'}\| - \frac{1}{m(m-1)} \sum_{j \neq j'} \|Y_j - Y_{j'}\|.$$

Under the null hypothesis that the two samples are drawn from the same distribution, the energy distance is zero. Statistical significance is assessed using a permutation procedure in which sample labels are randomly reassigned and the test statistic is recomputed for each permutation. The resulting empirical distribution is used to estimate a $p$-value in an analogous manner to other permutation-based tests.

The Energy Distance test is consistent against all fixed alternatives and is applicable in arbitrary dimensions, making it a powerful tool for detecting distributional differences in high-dimensional settings. Its nonparametric nature and reliance on pairwise distances align well with simulation-based and likelihood-free inference frameworks.

#### Maximum mean discrepancy test

The Maximum Mean Discrepancy (MMD) is a nonparametric statistical test used to determine whether two samples are drawn from the same underlying probability distribution. It is a kernel-based measure of discrepancy between distributions and is particularly effective for detecting differences in complex, high-dimensional data. The MMD test does not rely on parametric assumptions and is well suited for simulation-based and likelihood-free inference settings.

Let $X = \{X_1, \ldots, X_n\}$ and $Y = \{Y_1, \ldots, Y_m\}$ denote two independent samples drawn from distributions $P$ and $Q$, respectively. Given a reproducing kernel Hilbert space $\mathcal{H}$ with associated positive-definite kernel $k(\cdot, \cdot)$, the population MMD is defined as

$$\mathrm{MMD}(P, Q) = \|\mathbb{E}_{X \sim P}[k(X, \cdot)] - \mathbb{E}_{Y \sim Q}[k(Y, \cdot)]\|_{\mathcal{H}}.$$

An equivalent squared formulation that is more convenient for computation is given by

$$\mathrm{MMD}^2(P, Q) = \mathbb{E}_{X,X'}[k(X, X')] + \mathbb{E}_{Y,Y'}[k(Y, Y')] - 2\,\mathbb{E}_{X,Y}[k(X, Y)],$$

where $X'$ and $Y'$ are independent copies of $X$ and $Y$, respectively.

An unbiased empirical estimator of the squared MMD is

$$\widehat{\mathrm{MMD}}^2 = \frac{1}{n(n-1)} \sum_{i \neq i'} k(X_i, X_{i'}) + \frac{1}{m(m-1)} \sum_{j \neq j'} k(Y_j, Y_{j'}) - \frac{2}{nm} \sum_{i=1}^{n} \sum_{j=1}^{m} k(X_i, Y_j).$$

Under the null hypothesis $P = Q$, the MMD equals zero when a characteristic kernel is used. Statistical significance is assessed via a permutation or bootstrap procedure, in which sample labels are randomly permuted and the test statistic is recomputed to generate an empirical null distribution and corresponding $p$-value.

The flexibility afforded by the choice of kernel allows the MMD test to capture a wide range of distributional differences, including higher-order moments and nonlinear dependencies. As a result, MMD has become a widely used discrepancy measure in two-sample testing, generative model evaluation, and approximate Bayesian computation frameworks.

#### Statistical evaluation of Crocker plot distributions for model comparison

To justify the use of Bayesian information criterion (BIC) in light of the results of Marin et al. [4], we assess whether the posterior predictive summaries generated by each model, in our case Crocker plots, are statistically distinguishable. If the samplings of Crocker plots for each model are found to be statistically distinguishable, then that means the differences between the results of the two models is great enough that the BIC method should be able to choose which model is optimal. If they are not indistinguishable, then the BIC method may not give accurate results because the results of the two models are too similar. For each row of Table 1, we considered the posterior densities (an example is shown in Fig 4) and generated 500 Crocker plots for both $\text{Model}_{\text{AL}}$ and $\text{Model}_{\text{DO}}$. Parameter combinations were sampled according to their posterior probabilities, with acceptance probabilities proportional to the posterior density, resulting in retention of only the highest-probability parameter sets (approximately the top 1%). This procedure ensures that the resulting Crocker plots reflect the dominant posterior behavior of each model.

To compare the resulting collections of Crocker plots, we applied three complementary nonparametric two-sample tests: PERMANOVA, the Energy Distance test, and the MMD test. These tests were selected because they are well suited for high-dimensional summary statistics represented as matrices and are designed to assess differences between entire distributions rather than individual scalar values. For each test, the null hypothesis is that the distributions of Crocker plots generated by $\text{Model}_{\text{AL}}$ and $\text{Model}_{\text{DO}}$ are the same.

While all three methods test for distributional differences, they emphasize distinct aspects of the data. PERMANOVA assesses differences in group centroids within a specified distance space and is primarily sensitive to changes in the overall structure of the summary statistics, though it may also reflect differences in within-group dispersion. The Energy Distance test measures discrepancies between distributions based on pairwise distances and is consistent against all fixed alternatives, making it sensitive to differences in location, scale, and distributional shape. In contrast, MMD compares distributions through kernel mean embeddings and, depending on the choice of kernel, can capture higher-order and nonlinear differences that may not be detected by purely distance-based measures. Together, these tests provide a robust assessment of distributional distinguishability across multiple complementary notions of difference.

The resulting test statistics and associated permutation-based $p$-values are reported in Supplementary Material Table S1. In all cases, the $p$-values are less than 0.001, providing strong evidence against the null hypothesis and indicating that the posterior predictive distributions of Crocker plots differ significantly between the two models. Importantly, these results do not quantify model quality directly but instead establish that the summary statistics used for inference are meaningfully distinct across models. This distinction is a necessary condition for principled model comparison and supports the subsequent use of BIC to score and compare models based on their posterior predictive behavior.

| Ground-Truth Parameters | | | Permanova | | Energy Distance | | Maximum Mean Discrepancy | |
|---|---|---|---|---|---|---|---|---|
| C | L | W | Stat Value | P-value | Stat Value | P-value | Stat Value | P-value |
| 1.8 | 0.4 | 0 | 67.2 | 0.001 | 818 | 2.42e-38 | 0.025 | 1.33e-45 |
| 0.7 | 2.5 | 0 | 43 | 0.001 | 135 | 1.25e-24 | 0.014 | 4.91e-26 |
| 2 | 0.1 | 0 | 162 | 0.001 | 1.06e+03 | 2.12e-67 | 0.04 | 4.42e-72 |
| 0.9 | 0.6 | 0 | 16.6 | 0.001 | 351 | 4.57e-16 | 0.01 | 2.4e-19 |
| 0.5 | 0.5 | 0 | 151 | 0.001 | 895 | 5.51e-51 | 0.028 | 8.35e-51 |
| 0.2 | 1.5 | 0 | 93.3 | 0.001 | 516 | 1.77e-47 | 0.028 | 6.7e-51 |
| 1.5 | 0.7 | 0 | 171 | 0.001 | 241 | 1.15e-76 | 0.044 | 2.58e-79 |
| 2.5 | 2.5 | 0 | 65 | 0.001 | 109 | 6.92e-56 | 0.032 | 3.13e-57 |
| 2 | 1.5 | 0 | 48.8 | 0.001 | 81.2 | 2.99e-44 | 0.025 | 5.06e-46 |
| 1.2 | 0.7 | 0 | 141 | 0.001 | 223 | 8.31e-70 | 0.041 | 1.31e-72 |
| 1.8 | 0.4 | 0.05 | 244 | 0.001 | 2.14e+03 | 1.74e-102 | 0.066 | 6.53e-118 |
| 0.7 | 2.5 | 0.05 | 255 | 0.001 | 618 | 1.02e-106 | 0.062 | 1.95e-109 |
| 2 | 0.1 | 0.05 | 16 | 0.001 | 310 | 1.63e-18 | 0.012 | 1.06e-22 |
| 0.9 | 0.6 | 0.05 | 293 | 0.001 | 2.47e+03 | 3.77e-105 | 0.067 | 8.87e-119 |
| 0.5 | 0.5 | 0.05 | 109 | 0.001 | 998 | 3.82e-65 | 0.049 | 1.62e-87 |
| 0.2 | 1.5 | 0.05 | 463 | 0.001 | 1.29e+03 | 5.99e-119 | 0.067 | 2.34e-119 |
| 1.5 | 0.7 | 0.05 | 195 | 0.001 | 1.5e+03 | 5.21e-129 | 0.091 | 2.78e-160 |
| 2.5 | 2.5 | 0.05 | 370 | 0.001 | 553 | 4.04e-172 | 0.099 | 4.34e-173 |
| 2 | 1.5 | 0.05 | 1.3e+03 | 0.001 | 1.83e+03 | 1.87e-191 | 0.11 | 7.15e-193 |
| 1.2 | 0.7 | 0.05 | 1.29e+03 | 0.001 | 3.46e+03 | 1.73e-175 | 0.104 | 5.51e-183 |

**Supplementary Material Table S1. Model$_{\mathbf{AL}}$ versus Model$_{\mathbf{DO}}$ Crocker plot distribution statistical difference results.** We ran 500 approximate Bayesian computation (ABC) simulations of the top 1% of parameter combinations from the original simulation and generated their corresponding Crocker plots. Those 500 Crocker plots each were then statistically compared using three statistical tests: PERMANOVA Energy Distance, and MMD. The test statistics and $p$-values for each test and each ground-truth parameter combination from Table 1 are shown.

### Supplementary Material Appendix 4. Nearest neighbor t-SNE visualizations.

To investigate the differences between the with and without alignment groups, we modified the original t-SNE plot in Fig 2 b and c. Instead of coloring each point by their color coordinated by its location in 3D space, we colored all points with W=0 red and all points with W=0.05 in blue (see Supplementary Material Figure S1). Extending it further, for each row of Table 1, we highlighted the point with the ground truth C,L,W parameter values in yellow and found its nearest neighbor in t-SNE space of the other W value.

Through this new visualization of the Crocker summaries for the two different models, we see that they occupy two distinct regions of space. This can give a user confidence that model selection is a possibility. If these two models heavily overlapped, then there might not be enough information to distinguish between the two models during the model selection process.

### Supplementary Material Appendix 5. Akaike information criterion.

Akaike information criterion (AIC) is a widely used criterion for model selection and comparison in statistical modeling and machine learning. Like BIC, it evaluates the trade-off between model complexity and goodness of fit. However, AIC is derived from an information-theoretic perspective, specifically through the minimization of Kullback–Leibler divergence, and is primarily oriented toward predictive accuracy.
The AIC is defined as:

$$\mathrm{AIC} = -2\ln(\mathcal{L}) + 2(k+1)$$

where $\mathcal{L}$ is the likelihood of the model given the data, and $k$ is the number of parameters in the model. The log-likelihood $\ln(\mathcal{L})$ is defined as:

$$\ln(\mathcal{L}) = -(n/2)\ln(2\pi) - (n/2)\ln(\sigma^2) - \frac{SSE}{2\sigma^2}$$

where $\sigma^2 = SSE/n$.
The penalty term $2(k+1)$ discourages overly complex models by penalizing the inclusion of additional parameters, though less strongly than the $(k+1)\ln(n)$ term used in BIC. In contrast to BIC, which is motivated by Bayesian evidence and is asymptotically consistent for model selection, AIC does not possess this consistency property and instead prioritizes predictive performance. For this reason, BIC is more aligned with the objectives of the present study. Nevertheless, AIC provides a complementary perspective, and its inclusion allows for additional comparison across model selection criteria.

### Supplementary Material Appendix 6. Case Analysis: When $\mathbf{Model_{AL}}$ outperforms $\mathbf{Model_{DO}}$.

In Table 1, there are two cases in which $\mathrm{Model_{AL}}$ yields lower sum of squared errors ($SSE$) and BIC values than $\mathrm{Model_{DO}}$, despite the data being generated using $\mathrm{Model_{DO}}$. In the remaining eight cases, $\mathrm{Model_{DO}}$ attains lower $SSE$ and BIC values, consistent with the ground truth model. Below, we examine the two instances in which $\mathrm{Model_{AL}}$ outperforms $\mathrm{Model_{DO}}$ under these criteria.
In the first instance, corresponding to ground truth parameters (C, L) $= (0.5, 0.5)$, the median parameter estimates for $\mathrm{Model_{AL}}$ are (C, L, W) $= (0.751, 0.173, 0.012)$, while the median estimates for $\mathrm{Model_{DO}}$ are (C, L) $= (0.645, 0.169)$. In the second instance, corresponding to ground truth parameters (C, L) $= (0.2, 1.5)$, the median parameter estimates for $\mathrm{Model_{AL}}$ are (C, L, W) $= (0.142, 2.02, 0.008)$, while those for $\mathrm{Model_{DO}}$ are (C, L) $= (0.202, 1.566)$. Thus, although $\mathrm{Model_{AL}}$ achieves lower $SSE$ and BIC values in these two cases, its parameter estimates do not more closely recover the ground truth parameters. Rather, $\mathrm{Model_{AL}}$ identifies alternative parameter combinations that provide a slightly better fit to these particular simulated data sets. Notably, the estimated values of $W$ remain close to zero, indicating that the improved fit does not require substantial alignment behavior. Because $\mathrm{Model_{AL}}$ approaches $\mathrm{Model_{DO}}$ as $W$ approaches zero, these results illustrate that the two models can be difficult

to distinguish when the inferred alignment strength is small. Thus, the *SSE* and BIC values remain useful measures of relative model fit, but should be interpreted together with the inferred parameter values when distinguishing between these nested models.

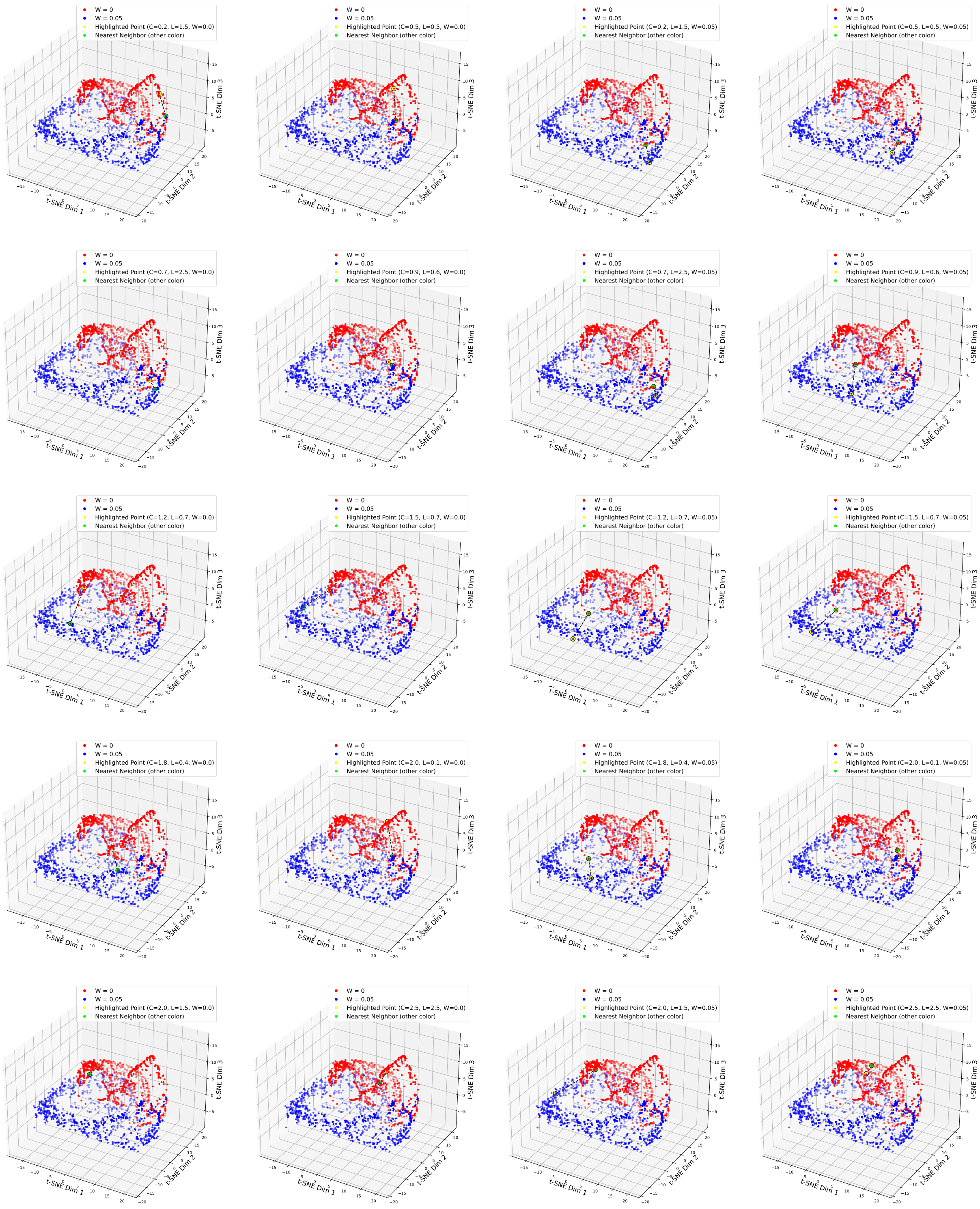


**Supplementary Material Figure S1.** T-SNE nearest-neighbor visualizations for the selected (C,L) pairs from Table 1 when W=0.0 (left two columns) and W=0.05 (right two columns). All points where W=0 are in red and similarly all points where W=0.05 are in blue. Each plot has one selected (C,L) pair at W=0.0 (highlighted in yellow) and its nearest neighbor where W=0.05 is highlighted in red for the left two columns and the reverse on the right (W=0.05 highlighted in yellow and nearest W=0.0 neighbor highlighted in red).

| Ground-Truth Parameters | | | $\text{Model}_{AL}$ | | | $\text{Model}_{DO}$ | | AIC | | |
|---|---|---|---|---|---|---|---|---|---|---|
| C | L | W | $\text{C}_{\text{estimate}}$ | $\text{L}_{\text{estimate}}$ | $\text{W}_{\text{estimate}}$ | $\text{C}_{\text{estimate}}$ | $\text{L}_{\text{estimate}}$ | $\text{Model}_{AL}$ | $\text{Model}_{DO}$ | $\text{Model}_{AL}$-$\text{Model}_{DO}$ |
| 1.8 | 0.4 | 0 | 2.32 | 0.311 | 0.005 | 1.825 | 0.394 | 2.22e+05 | 2.00e+05 | 2.19e+04 |
| 0.7 | 2.5 | 0 | 0.687 | 2.398 | 0.004 | 0.669 | 2.321 | 2.21e+05 | 2.07e+05 | 1.37e+04 |
| 2.0 | 0.1 | 0 | 1.351 | 0.149 | 0.003 | 1.404 | 0.123 | 2.48e+05 | 2.44e+05 | 4.79e+03 |
| 0.9 | 0.6 | 0 | 0.961 | 0.315 | 0.029 | 0.888 | 0.592 | 3.10e+05 | 2.22e+05 | 8.81e+04 |
| 0.5 | 0.5 | 0 | 0.751 | 0.173 | 0.012 | 0.645 | 0.169 | 3.60e+05 | 3.65e+05 | -5.23e+03 |
| 0.2 | 1.5 | 0 | 0.142 | 2.020 | 0.008 | 0.202 | 1.566 | 2.76e+05 | 2.80e+05 | -3.61e+03 |
| 1.5 | 0.7 | 0 | 2.107 | 0.661 | 0.004 | 1.553 | 0.705 | 2.22e+05 | 1.83e+05 | 3.86e+04 |
| 2.5 | 2.5 | 0 | 2.391 | 2.333 | 0.002 | 2.558 | 2.506 | 1.86e+05 | 1.82e+05 | 4.38e+03 |
| 2.0 | 1.5 | 0 | 2.226 | 1.699 | 0.002 | 2.000 | 1.499 | 1.88e+05 | 1.56e+05 | 3.25e+04 |
| 1.2 | 0.7 | 0 | 2.233 | 0.698 | 0.009 | 1.443 | 0.632 | 2.73e+05 | 2.00e+05 | 7.27e+04 |
| 1.8 | 0.4 | 0.05 | 2.064 | 0.351 | 0.058 | 1.583 | 0.511 | 2.45e+05 | 3.12e+05 | -6.61e+04 |
| 0.7 | 2.5 | 0.05 | 0.703 | 2.564 | 0.049 | 0.501 | 1.227 | 2.01e+05 | 3.29e+05 | -1.28e+05 |
| 2.0 | 0.1 | 0.05 | 1.389 | 0.146 | 0.047 | 0.794 | 0.591 | 2.55e+05 | 3.17e+05 | -6.19e+04 |
| 0.9 | 0.6 | 0.05 | 1.051 | 0.517 | 0.058 | 1.625 | 0.492 | 2.14e+05 | 3.47e+05 | -1.33e+05 |
| 0.5 | 0.5 | 0.05 | 0.469 | 0.416 | 0.077 | 0.756 | 0.676 | 2.77e+05 | 3.71e+05 | -9.39e+04 |
| 0.2 | 1.5 | 0.05 | 0.214 | 1.522 | 0.053 | 0.525 | 0.913 | 2.42e+05 | 3.23e+05 | -8.08e+04 |
| 1.5 | 0.7 | 0.05 | 1.612 | 0.747 | 0.050 | 1.867 | 0.499 | 1.90e+05 | 2.52e+05 | -6.22e+04 |
| 2.5 | 2.5 | 0.05 | 2.605 | 2.372 | 0.051 | 1.882 | 0.514 | 1.85e+05 | 2.49e+05 | -6.41e+04 |
| 2.0 | 1.5 | 0.05 | 2.091 | 1.417 | 0.065 | 1.920 | 0.494 | 1.91e+05 | 3.00e+05 | -1.09e+05 |
| 1.2 | 0.7 | 0.05 | 1.208 | 0.694 | 0.068 | 1.894 | 0.486 | 1.86e+05 | 2.18e+05 | -3.22e+04 |

**Supplementary Material Table S2. Model selection comparison results of $\textbf{Model}_{\textbf{AL}}$ to $\textbf{Model}_{\textbf{DO}}$ using AIC.** Similar to Table 1, results are summarized using AABC-estimated parameter values for both models, the AIC score and sum of squared errors ($SSE$) for each model, and the difference of AIC scores. This is done for the ten ground truth samples chosen in (d) and (e) of Figure 2 corresponding to the selection of C, L, and W values in the first three columns.

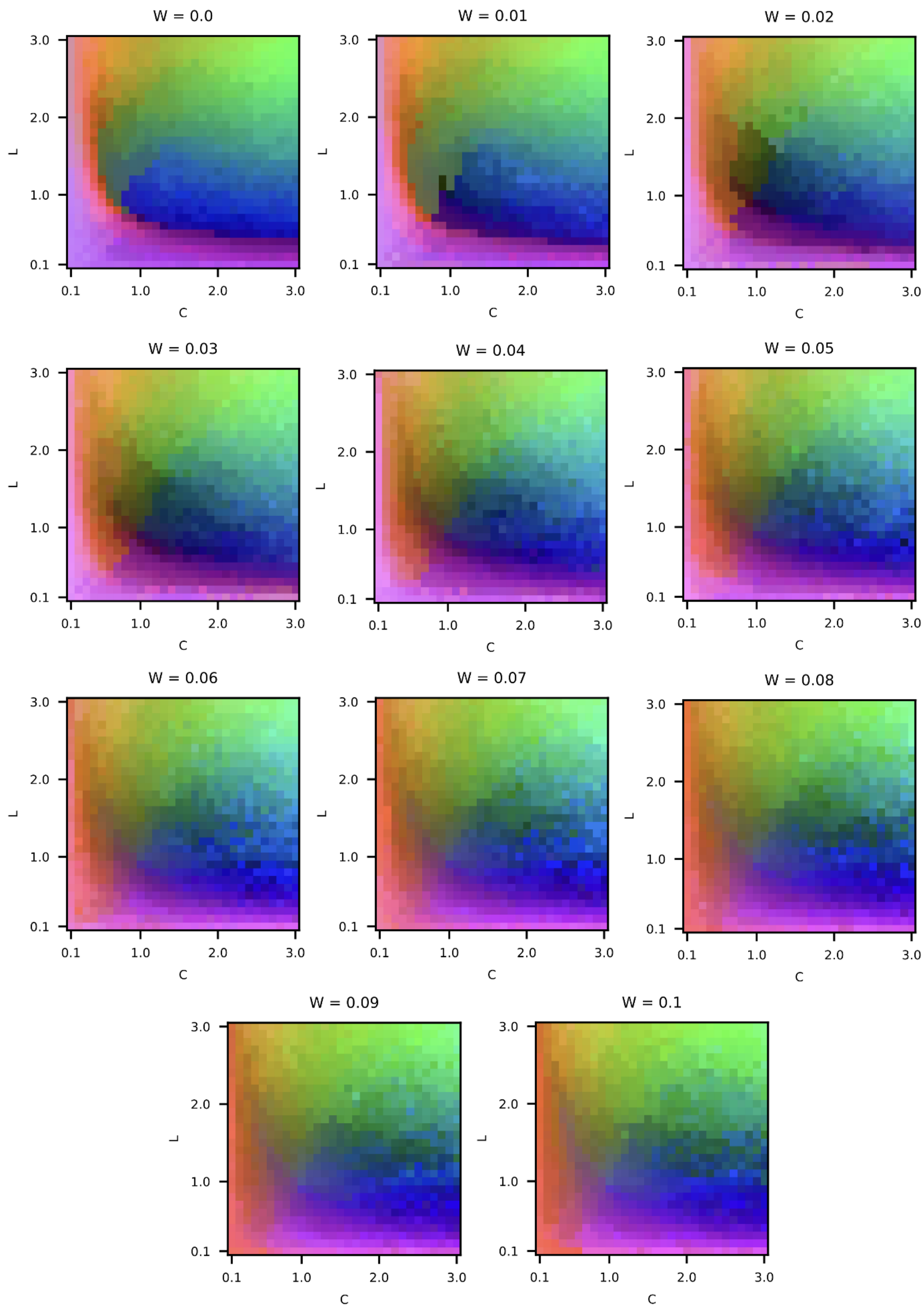


**Supplementary Material Figure S2.** The full selection of the Betti-0 and Betti-1 t-SNE colormaps, showcasing an example of no alignment (W=0.0, top left) and different levels of alignment (W=0.01-W=0.1). The colors for each C, L, W combination are the colors that were generated in the 3D t-SNE plot ((c) of Fig 2) for the corresponding C, L, W point.

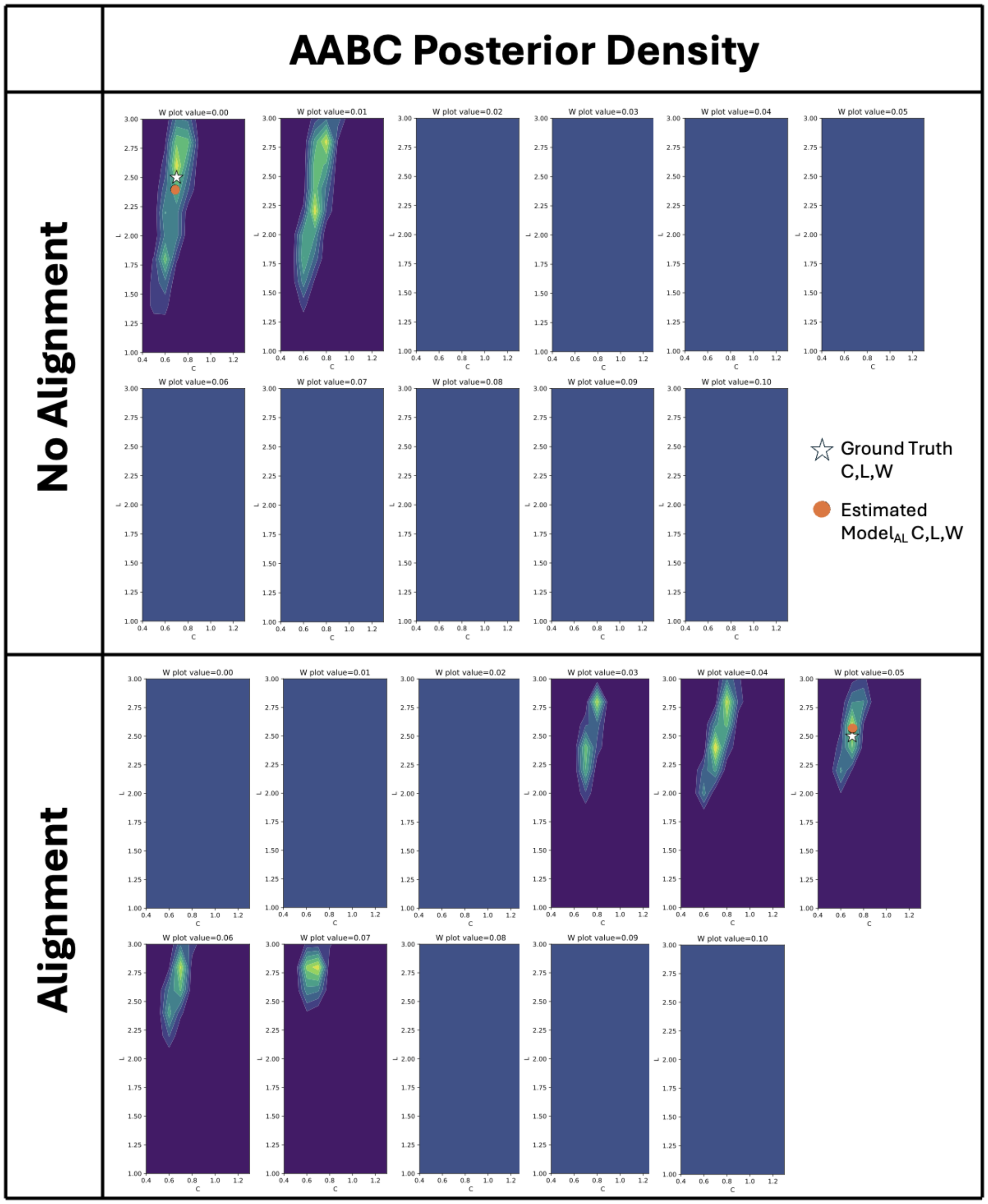


**Supplementary Material Figure S3.** This is the full selection of approximate approximate Bayesian computation (AABC) posterior density plots for Model$_{\mathrm{AL}}$. Top and bottom use data generated with no alignment (W=0.0) and alignment (W=0.05), respectively. The white star represents the ground-truth values whereas the orange dots represent the sample median AABC-estimated values for Model$_{\mathrm{AL}}$.

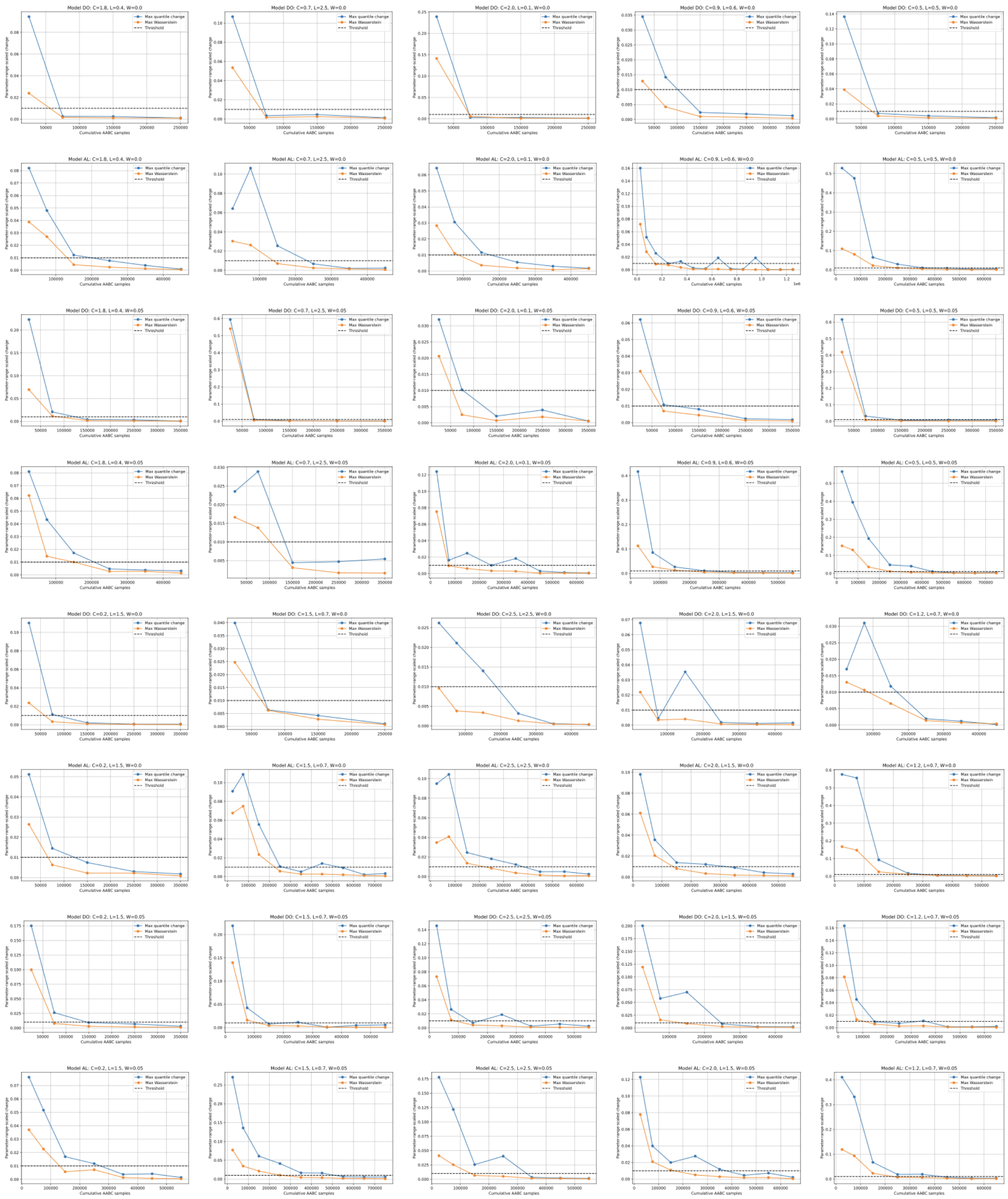


**Supplementary Material Figure S4.** The full selection of convergence plots for each row of Table 1 for both $\text{Model}_{\text{DO}}$ and $\text{Model}_{\text{AL}}$ when the ground truth value for W is 0 and 0.05 for a total of four convergence plots per row. AABC samples were generated iteratively until both the maximum normalized posterior-quantile change and the maximum normalized Wasserstein distance were less than 0.01 for three consecutive checkpoint comparisons.
