## Supplementary figures and images for "Topologically-based parameter inference for agent-based model selection from spatiotemporal cellular data"

### Supplementary Video 1

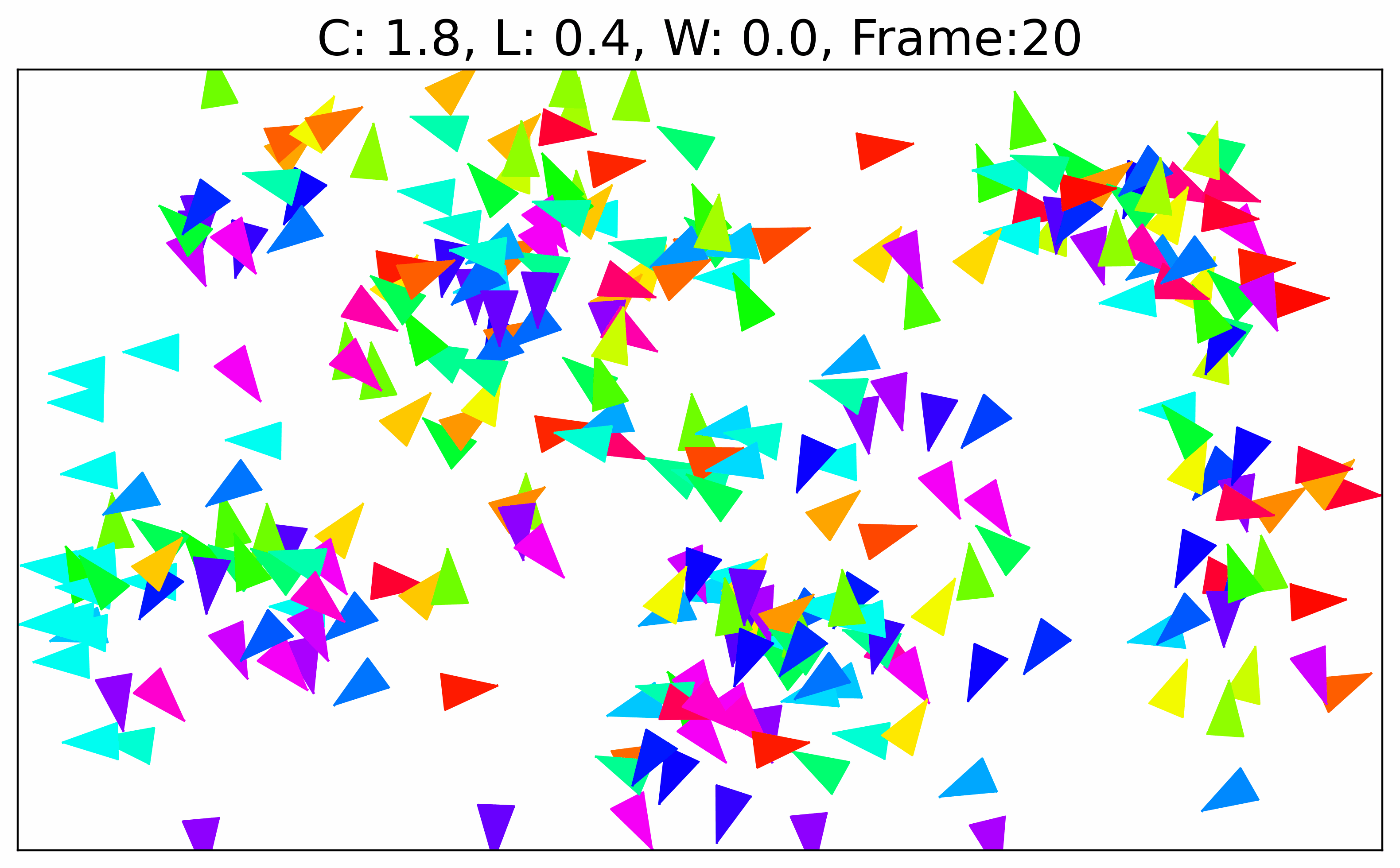

### Supplementary Video 2

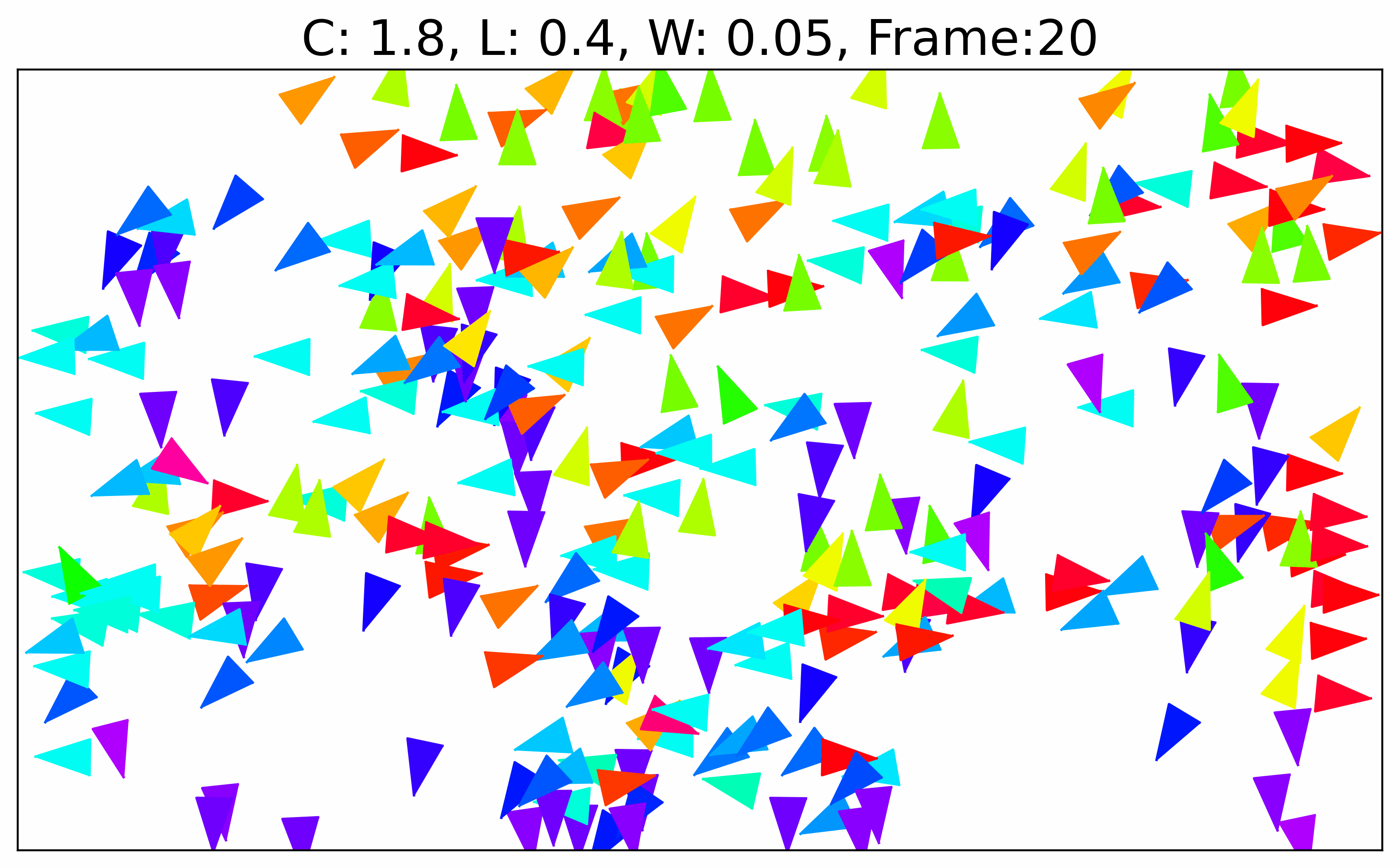

### Supplementary Video 3

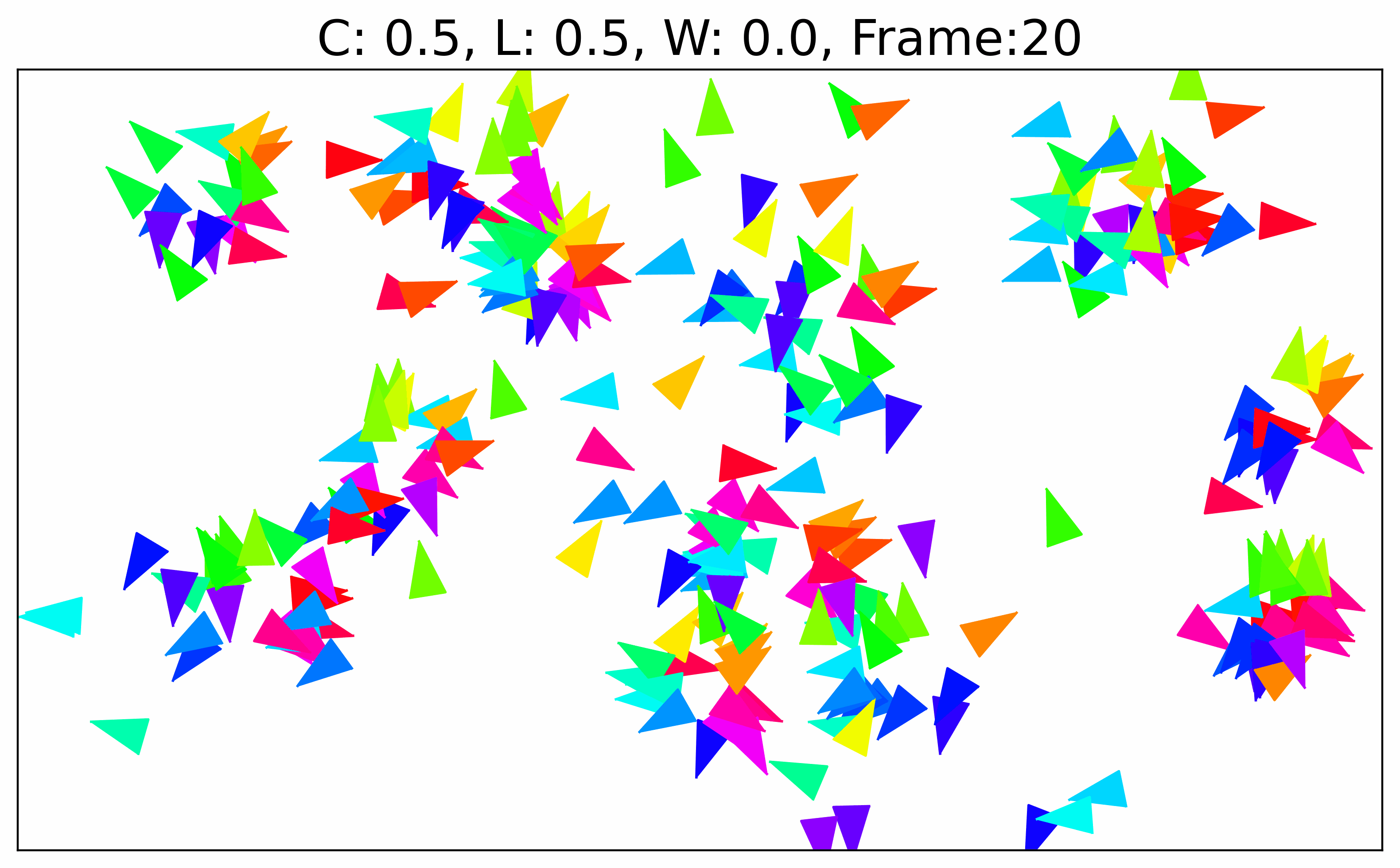

### Supplementary Video 4

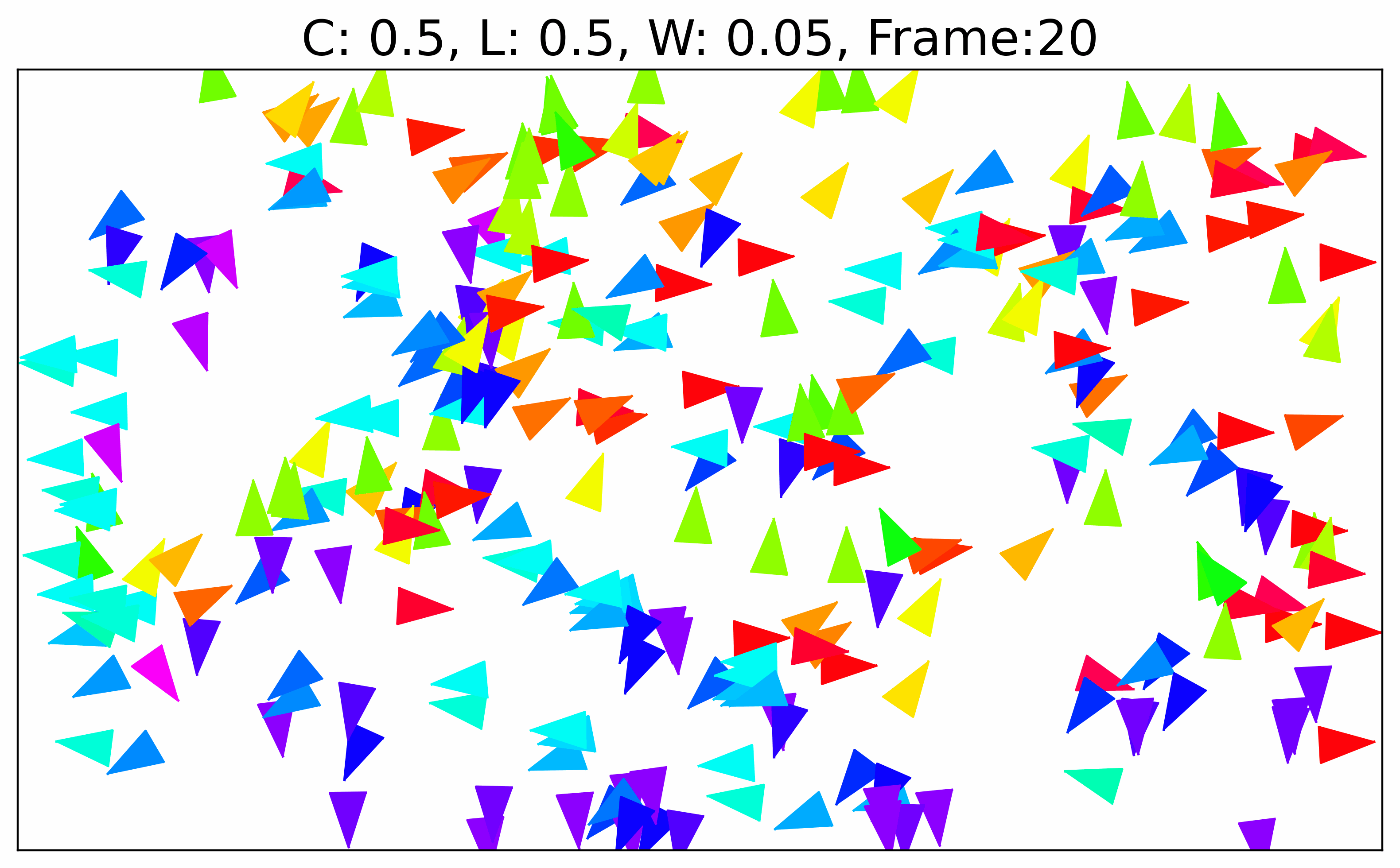
